# *De novo* design of high-affinity antibody variable regions (Fv) against the SARS-CoV-2 spike protein

**DOI:** 10.1101/2020.04.09.034868

**Authors:** Veda Sheersh Boorla, Ratul Chowdhury, Costas D. Maranas

## Abstract

The emergence of SARS-CoV-2 is responsible for the pandemic of respiratory disease known as COVID-19, which emerged in the city of Wuhan, Hubei province, China in late 2019. Both vaccines and targeted therapeutics for treatment of this disease are currently lacking. Viral entry requires binding of the viral spike receptor binding domain (RBD) with the human angiotensin converting enzyme (hACE2). In an earlier paper^1^, we report on the specific residue interactions underpinning this event. Here we report on the *de novo* computational design of high affinity antibody variable regions through the recombination of VDJ genes targeting the most solvent-exposed hACE2-binding residues of the SARS-CoV-2 spike protein using the software tool OptMAVEn-2.0^2^. Subsequently, we carry out computational affinity maturation of the designed prototype variable regions through point mutations for improved binding with the target epitope. Immunogenicity was restricted by preferring designs that match sequences from a 9-mer library of “human antibodies” based on H-score (human string content, HSC)^3^. We generated 106 different designs and report in detail on the top five that trade-off the greatest affinity for the spike RBD epitope (quantified using the Rosetta binding energies) with low H-scores. By grafting the designed Heavy (VH) and Light (VL) chain variable regions onto a human framework (Fc), high-affinity and potentially neutralizing full-length monoclonal antibodies (mAb) can be constructed. Having a potent antibody that can recognize the viral spike protein with high affinity would be enabling for both the design of sensitive SARS-CoV-2 detection devices and for their deployment as therapeutic antibodies.

## Main

Over the last few weeks, several studies on using human or humanized antibodies targeted at the SARS-CoV-2 spike protein have been reported^4,5,6,7^. In addition, multiple efforts by laboratories and companies (Cellex, GeneTex etc.) for the development of antibody-based tests for SARS-CoV-2 detection are ongoing^8^. At the same time, significant progress towards the isolation and design of vaccines (mRNA-1273 vaccine © 2020 Moderna, Inc) and neutralizing antibodies^9^ has been made. A computational study identified the structural basis for multi-epitope vaccines^10,11^ whereas in another study, the glycosylation patterns of the spike SARS-CoV-2 protein were computationally deduced^12^. In addition, fully human single domain anti-SARS-CoV-2 antibodies with sub-nanomolar affinities were identified from a phage-displayed single-domain antibody library. Naïve CDRs were grafted to framework regions of an identified human germline IGHV allele using SARS-CoV-2 RBD and S1 protein as antigens^13^. In another study^14^, a human antibody 47D11 was identified to have cross neutralizing effect on SARS-CoV-2 by screening a library of SARS-CoV-1 antibodies. In two other studies, potent neutralizing antibodies were isolated from the sera of convalescent COVID-19 patients^15,16^. To the best of our knowledge, none of these neutralizing antibody sequences are publicly available. In a follow up effort^17^, human antibody CR3022 (which is neutralizing against SARS-CoV-1^18^) has been shown to bind to SARS-CoV-2 RBD in a cryptic epitope but without a neutralizing effect for SARS-CoV-2 *in vitro*. Desautels et al^19^, performed a machine learning based *in silico* mutagenesis of SARS-CoV-1 neutralizing antibodies to bind to SARS-CoV-2 spike protein. Walter et al^20^ generated a number of synthetic nanobodies by *in vitro* screening large combinatorial libraries for binding to SARS-CoV-2 spike RBD^20^. However, studies that perform structure guided design of high affinity antibodies against specific epitopes of SARS-CoV-2 spike protein that may interfere with hACE2 binding are still lacking.

Motivated by these shortcomings, here we explore the *de novo* design of antibody variable regions targeting the most solvent-exposed residues of the spike protein that are also part of the residue contact map involved in hACE2 binding, and trade-off binding energy against human sequence content in the variable region. The goal here is to exhaustively explore the sequence space of all possible variable region designs and report a range of diverse solutions that can serve as potentially neutralizing antibodies (nAb). We find that many different combinations of VDJ genes followed by mutation can yield potentially high affinity variable regions (scored using the Rosetta binding energy function) against an epitope of the spike protein RBD. Pareto optimal designs with respect to binding affinity vs. human content were drawn and five affinity matured designs are detailed in the results section.

We first performed solvent accessibility analysis using the STRIDE^21^ program on the 21 hACE2-binding residues of the SARS-CoV-2 spike protein (S-protein) RBD to define our binding epitope. The top seven residues with the highest solvent accessibility scores (i.e., SAS) are (Arg346, Phe347, Ala348, Tyr351, Ala352, Asn354, and Arg355) comprising our binding epitope (see Figure 1). Furthermore, the epitope is accessible for binding to RBD in the open confirmation of the full spike protein (See Supp. Fig. S8).

**Figure 1.**
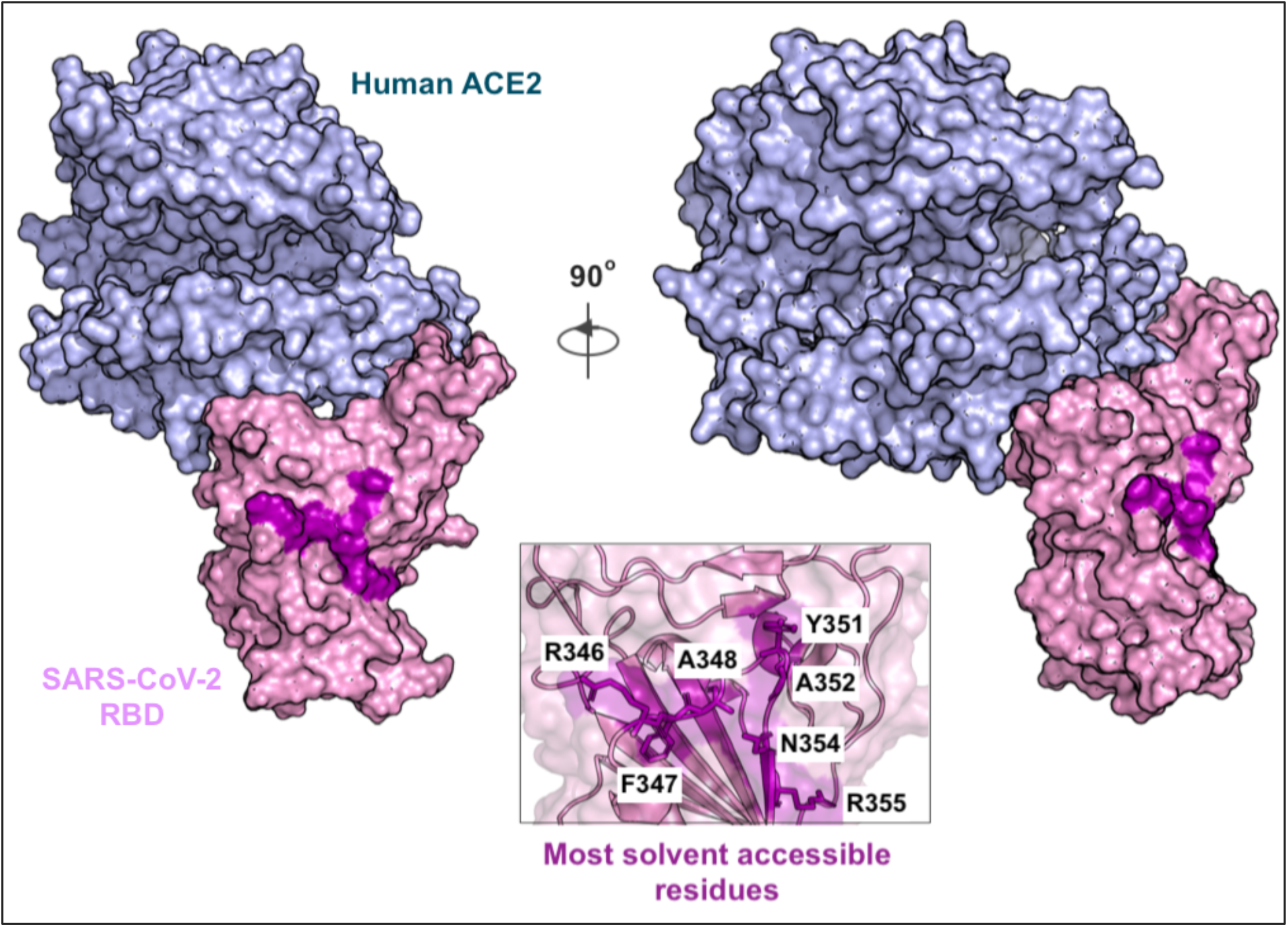
The SARS-CoV-2 spike protein RBD in complex with Human ACE2 protein (PDB-id: 6LZG) is shown along with the most solvent accessible residues at the binding interface highlighted in purple. A zoomed view of these seven epitope residues is shown in the inset box. The numbering scheme for the S-protein residues is same as in PDB accession id 6LZG (rcsb.org/structure/6LZG or 6VW1^7^ and 6M0J^6^).

We next used the previously developed OptMAVEn-2.0^2^ software to computationally identify the combination of VDJ genes forming the variable region that best binds the desired epitope. OptMAVEn^22^ has been used before successfully to design five high affinity CDRs against a FLAG tetrapeptide^23^, three thermally and conformationally stable antibody variable regions (sharing less than 75% sequence similarity to any naturally occurring antibody sequence) against a dodecapeptide mimic of carbohydrates^24^ and two thermostable, high affinity variable heavy chain domains (V_H_H) against α-synuclein peptide responsible for Parkinson’s disease^25^. All these designs were experimentally constructed and nanomolar affinities for their respective target antigens was demonstrated.

Through a combination of rotations and translations, OptMAVEn-2.0 identified 3,234 unique antigen poses that presented the epitope to the antibody differently. The combinatorial space of different VDJ genes that upon recombination form the variable region of the prototype antibody was informed by the MAPs database of antibody parts^26^. MAPs (see Supp. Info. S1 for link to full database) contains 929 modular antibody (i.e., variable-V*, complementarity determining -CDR3, and joining-J*) parts from 1,168 human, humanized, chimeric, and mouse antibody structures (last updated in 2013). MAPs follows the antibody parts residue numbering convention as listed in the International iMmunoGeneTics (IMGT)^27^ database. IMGT catalogs antibody parts as variable (V), diversity (D) and joining (J) structure libraries. MAPs stores complete CDR3 parts, C-terminus-shortened V parts (i.e. V* parts) and N-terminus-shortened J parts (J* parts). Note that CDR3 includes the entire D gene and also up to the C-terminus of the V gene and up to the N-terminus of the J gene. In the remainder of the manuscript, the list of parts used to design the variable region are referred to as CDR3, V* and J* parts.

For each one of the 3,234 spike poses, OptMAVEn-2.0 identified a variable region combination composed of end-to-end joined V*, CDR3, and J* region parts that minimized the Rosetta binding energy between the variable region and spike epitope formed by the seven residues. As part of OptMAVEn-2.0, the combinatorial optimization problem was posed and solved as a mixed-integer linear programming (MILP) problem using the cplex solver^28^. The solution of this problem identifies, for each one of the spike poses, the complete design of the variable region using parts denoted as HV*, HCDR3, HJ* for the heavy chain H and L/KV*, L/KCDR3 and L/KJ* for the light chain-L/K. Only 173 antigen-presenting poses out of 3,234 explored, yielded non-clashing antigen-antibody complexes. These 173 poses were ranked on the basis of their Rosetta binding energies with the spike epitope and classified into 27 clusters (using *k-means*^29^) in a 19-dimensional space defined by quantitative descriptors of sequence similarity, three-dimensional spatial pose, and the angle at which they bind to the target epitope (see details in original paper^2^). The top five prototype designs with the highest Rosetta binding energies were present in four clusters and spanned a highly diverse set of choices of MAPs parts (see Table 1) with minimal conservation of the same part among the five prototype designs. The number entries in Table 1 correspond to the id of the gene in the MAPs database (which are identical to the ids used in IMGT). Note that design P5 uses a lambda (L) light chain instead of a kappa (K). Figure 1a plots the pairwise sequence similarity scores of the five antibody variable domains that were used in the top five designs. As expected, the top five prototype designs P1, P2, P3, P4, and P5 are the most dissimilar in their respective CDR3 domains in both light L, heavy H and HV* domain (but not LV*). They are the most similar in the choice of parts for the J* domains (see Figure 2a) reflecting the lack of diversity among possible choices for the J* domains in the MAPs database.

**Table 1.**
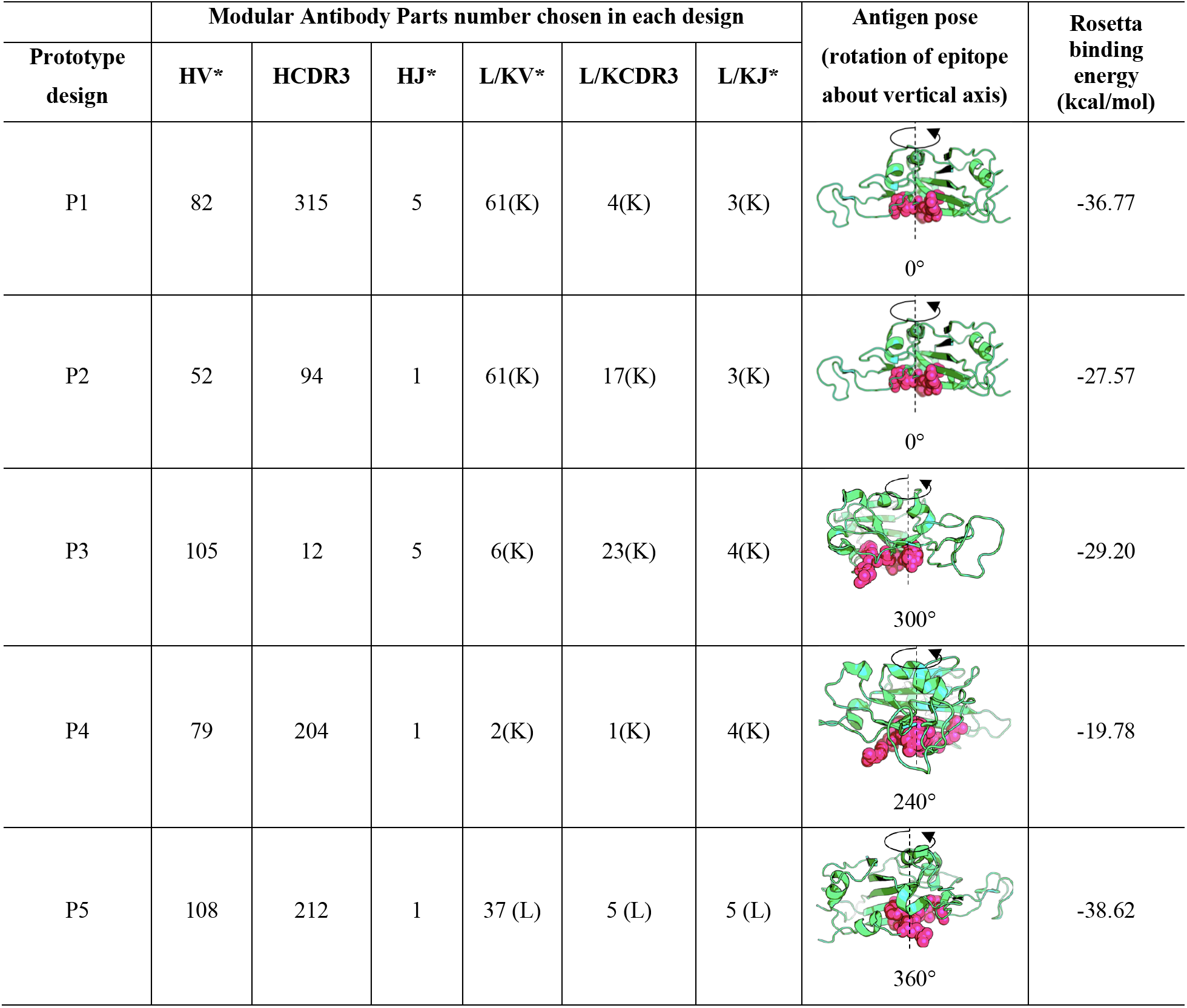
V*, CDR3, J* gene ids for the top five prototype variable region designs and corresponding Rosetta binding energies^30,31^.Antigen poses are described with the angle that the vertical axis through the epitope (shown in pink) centroid and the Cβ carbon of the residue with greatest z-axis coordinate forms.

| Prototype design | Modular Antibody Parts number chosen in each design |  |  |  |  |  | Antigen pose (rotation of epitope about vertical axis) | Rosetta binding energy (kcal/mol) |
| --- | --- | --- | --- | --- | --- | --- | --- | --- |
|  | HV* | HCDR3 | HJ* | L/KV* | L/KCDR3 | L/KJ* |  |  |
| P1 | 82 | 315 | 5 | 61(K) | 4(K) | 3(K) | 0° | -36.77 |
| P2 | 52 | 94 | 1 | 61(K) | 17(K) | 3(K) | 0° | -27.57 |
| P3 | 105 | 12 | 5 | 6(K) | 23(K) | 4(K) | 300° | -29.20 |
| P4 | 79 | 204 | 1 | 2(K) | 1(K) | 4(K) | 240° | -19.78 |
| P5 | 108 | 212 | 1 | 37 (L) | 5 (L) | 5 (L) | 360° | -38.62 |

**Figure 2.**
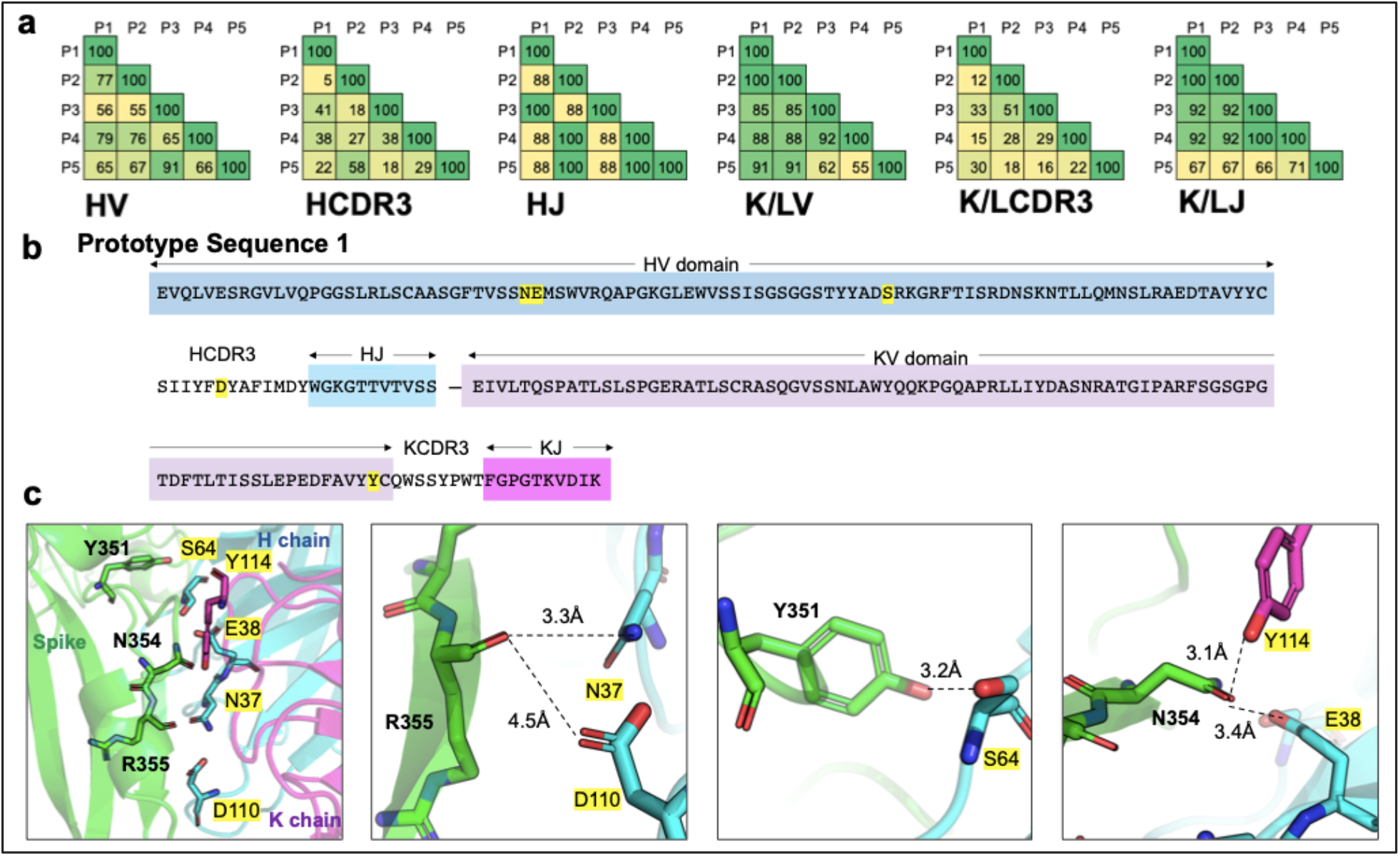
**(a)** Pairwise sequence similarity percentages between the members of the six parts that were used to construct the top five prototype variable regions with the lowest Rosetta binding energies with the viral spike epitope. **(b)** The amino acid sequence of prototype design P1 with the different domain parts highlighted in different colors. Spike epitope binding residues are highlighted in yellow. **(c)** Structural view of the strongest epitope-prototype variable region interactions for P1. They imply strong electrostatic capture of three epitope residues by five variable region residues spanning both heavy (H) and light (K) chains.

Inspection of the interaction of design P1 with the spike epitope reveals strong electrostatic contacts between the S-protein residues Tyr351, Asn354, and Arg355 (see Figure 1c) all of which have been deemed important for hACE2 binding^1^. The strongest contacts with the three epitope residues are established by five antibody residues spanning both the heavy and light chains (shown in yellow in Figure 2b). Spike Tyr351 interacts with Ser64 in the HV* domain, Asn354 interacts with Glu38 and Tyr114 in HV* and KV* domains respectively, while spike Arg355 interacts with Asn37 and Asp110 of HV* and HCDR3 domains, respectively, in the stable spike-antibody complex (see Figure 2c).

We next applied Rosetta-based *in silico* affinity maturation (see Methods) for each one of the top five prototype designs shown in Table 1 to further enhance the non-covalent binding between the antibody variable domains and the SARS-CoV-2 spike RBD. This computationally mimics the process of somatic hypermutation leading to eventual affinity maturation of antibodies in B cells. This procedure identified a total of 124 unique variable designs by introducing mutations in the five prototypes (see Figure 3a). We retained 106 designs which achieved both an improvement in the Rosetta binding energy over their respective prototype sequences and also further stabilization (i.e., lower overall Rosetta energy) of the spike-antibody complex (see upper right quadrant of Figure 3a). On average, upon affinity maturation, the binding energy was improved by ~14 kcal/mol and the overall energy was improved by ~37 kcal/mol. Supplementary S2 lists first the starting prototype design (i.e., P1, P2, P3, P4 or P5) followed by the 106 affinity matured designs (labeled as P1.D1, P1.D2, etc). On average, there were 4.5 mutations (Supp. info. S3) between computational affinity matured and prototype variable region designs.

**Figure 3.**
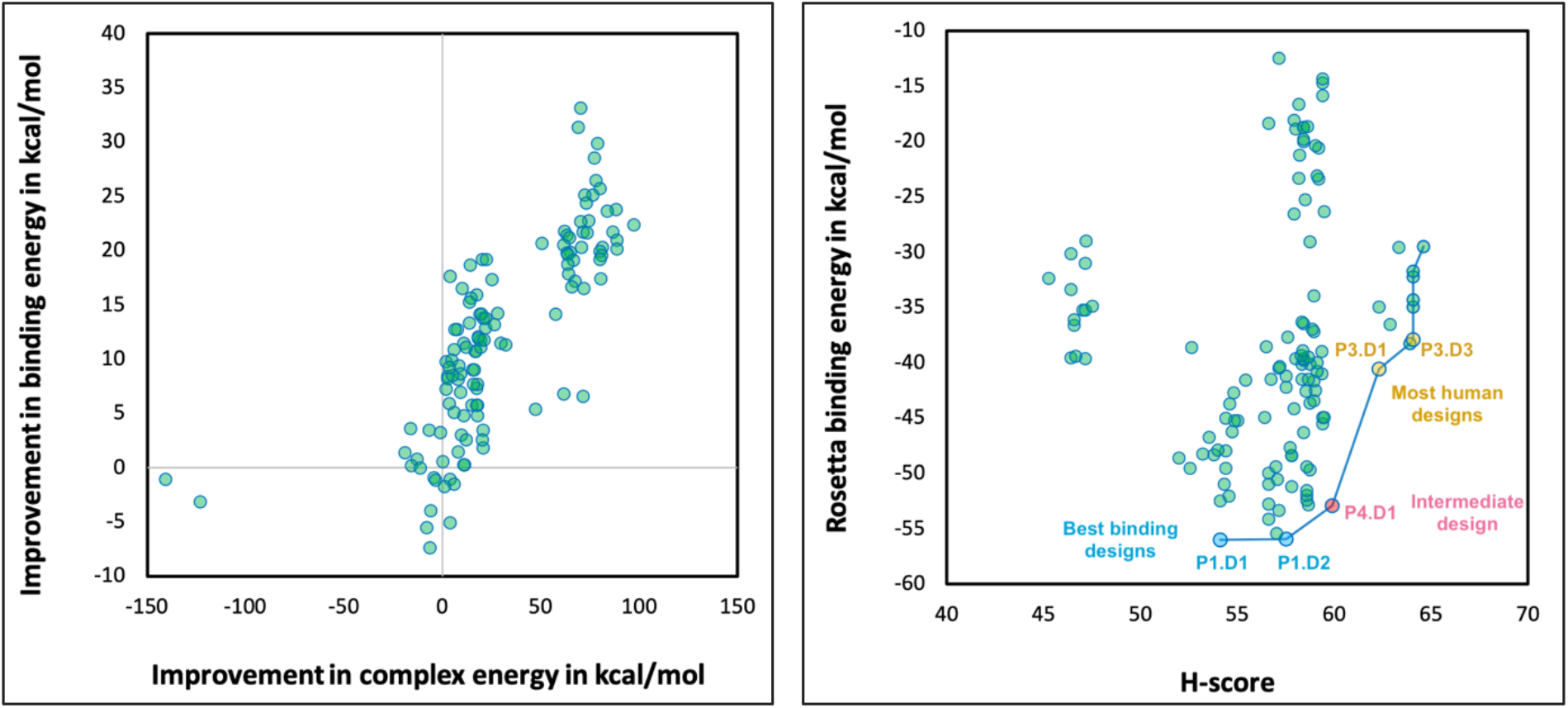
**(a)** The 106 out of 124 Rosetta-affinity matured designs that improve upon both energy criteria fall in the top right quadrant. **(b)** Plot of the Rosetta binding energy vs. H-score for the 106 retained affinity-matured sequences. The blue line connects the designs on the Pareto optimum curve between these two design objectives. The three best-binding affinity matured designs (emerging from P1) – P1.D1 and P1.D2 are shown in blue whereas the two most human designs (emerging from P3) – P3.D1 and P3.D3 are shown in yellow. An intermediate design P4.D1 (emerging from P4) is shown in red.

We next assessed the departure of the 106 designed variable regions from fully-human antibody sequences using H-Score^3^. H-score is defined as the sum of the minimum edit distance between all possible 9-mer fragments in the designed variable region from a library of all 9-mer sequences observed in human antibodies^22^. The value of H-score is scaled to 100 and normalized by the length of the sequence. An antibody sequence with all 9-mers exactly matching 9-mers of human antibodies will have a perfect H-score of 100. Figure 3b illustrates the trade-off between the Rosetta binding energy vs. H-score for these affinity matured variable region designs. For comparison, we calculated the H-score for the human antibodies CR3022^32^, 80R^33^, S230^34^ and M396^35^ which are known to be neutralizing against SARS-CoV-1. These antibodies (including only Fv regions) have an average H-score of 62.98 (stdev: 4.9) which are in the same range as our most human designs (e.g., P3.D1 and P3.D3).

We selected five designs that were on the Pareto optimum curve shown in Figure 3b. The Pareto optimum curve is defined as the collection of designs for which no other design exists that can simultaneously improve upon both criteria (i.e., Rosetta binding energy and H-score). Designs P1.D1 and P1.D2 shown in blue (in Fig. 3b) have the lowest Rosetta binding energies whereas P3.D1 and P3.D3 shown in yellow correspond to the ones with the highest H-scores. Design P4.D1 is an intermediate design that balances both binding energy and H-score. The lowest binding energy designs (P1.D1, P1.D2), irrespective of H-score, would be relevant in ELISA-based *in vitro* detection assays whereas the lowest H-score designs (P3.D1, P3.D3) may offer the highest potential as therapeutic antibodies. In addition, we calculated the Rosetta binding energy between the human CR3022^17^ (anti-SARS-CoV-1 antibody) and the SARS-CoV-2 spike protein RBD using complex structure (PDB-id:6W41) to be −56.4 kcal/mol which is very close to the Rosetta binding energy of designs P1.D1 and P1.D2. However, P1.D1 and P1.D2 bind a different epitope on the spike RBD than the one that CR3022 targets (see Supp. Fig S7).

Figure 4 shows the sequence alignment of these five selected affinity matured sequences (i.e., P1.D1, P1.D2, P4.D2, P3.D1, and P3.D3). Shown in red boxes are the conserved positions and in red font the positions with different but of similar type residues. A total of 156 out of 226 aligned positions are conserved among all designs. Table 2 lists the most important (strongest) contacts with the spike protein as informed by an *in silico* alanine scanning (Supp. info. S4) on the spike-binding residues of the variable region designs. In essence, the alanine scanning analysis identifies the loss in binding energy that is incurred upon mutating each residue (one at a time) to alanine.

**Table 2.** List of important contacts between the spike protein epitope residues and residues of each of the selected affinity matured designs. For each contact, the loss in binding energy upon mutation of antibody residue from the interface to alanine is tabulated. The corresponding interacting spike residue is also shown.

| Matured antibody Design id | Interface residue from antibody | Interacting spike residue(s) | Loss in binding energy upon mutation to alanine (kcal/mol) | Corresponding variable region |
| --- | --- | --- | --- | --- |
| P1.D1 | G62 | A352 | 9.69 | HV* |
|  | G63 | Y351 | 2.13 | HV* |
|  | L65 | Y489 | 1.26 | HV* |
|  | Y66 | S349 | 0.53 | HV* |
|  | I56 | C488 | 0.94 | HV* |
| P1.D2 | F109 | K356 | 2.77 | HCDR3 |
|  | T65 | Y351 | 1.42 | HV* |
|  | Y66 | A348,S349 | 1.06 | HV* |
|  | I56 | C488 | 0.94 | HV* |
|  | S57 | A352 | 0.71 | HV* |
| P4.D1 | A56 | L452 | 0.573 | KV* |
|  | D35 | R357 | 0.08 | HV* |
|  | G28 | T478,G482 | 0.01 | KV* |
|  | T85 | N481 | 0.00 | KV* |
|  | L67 | V445 | 0.00 | KV* |
| P3.D1 | W64 | A352 | 2.30 | HV* |
|  | N57 | N354 | 0.65 | HV* |
|  | F107 | T345 | 0.57 | KCDR3 |
|  | S29 | E340 | 0.14 | HV* |
|  | S108 | R346 | 0.12 | KCDR3 |
| P3.D3 | N57 | N354 | 0.99 | HV* |
|  | F107 | T345 | 0.33 | KCDR3 |
|  | D38 | R346 | 0.18 | KV* |
|  | S29 | E340 | 0.22 | HV* |
|  | Q106 | R346 | 0.03 | KCDR3 |

**Figure 4.**
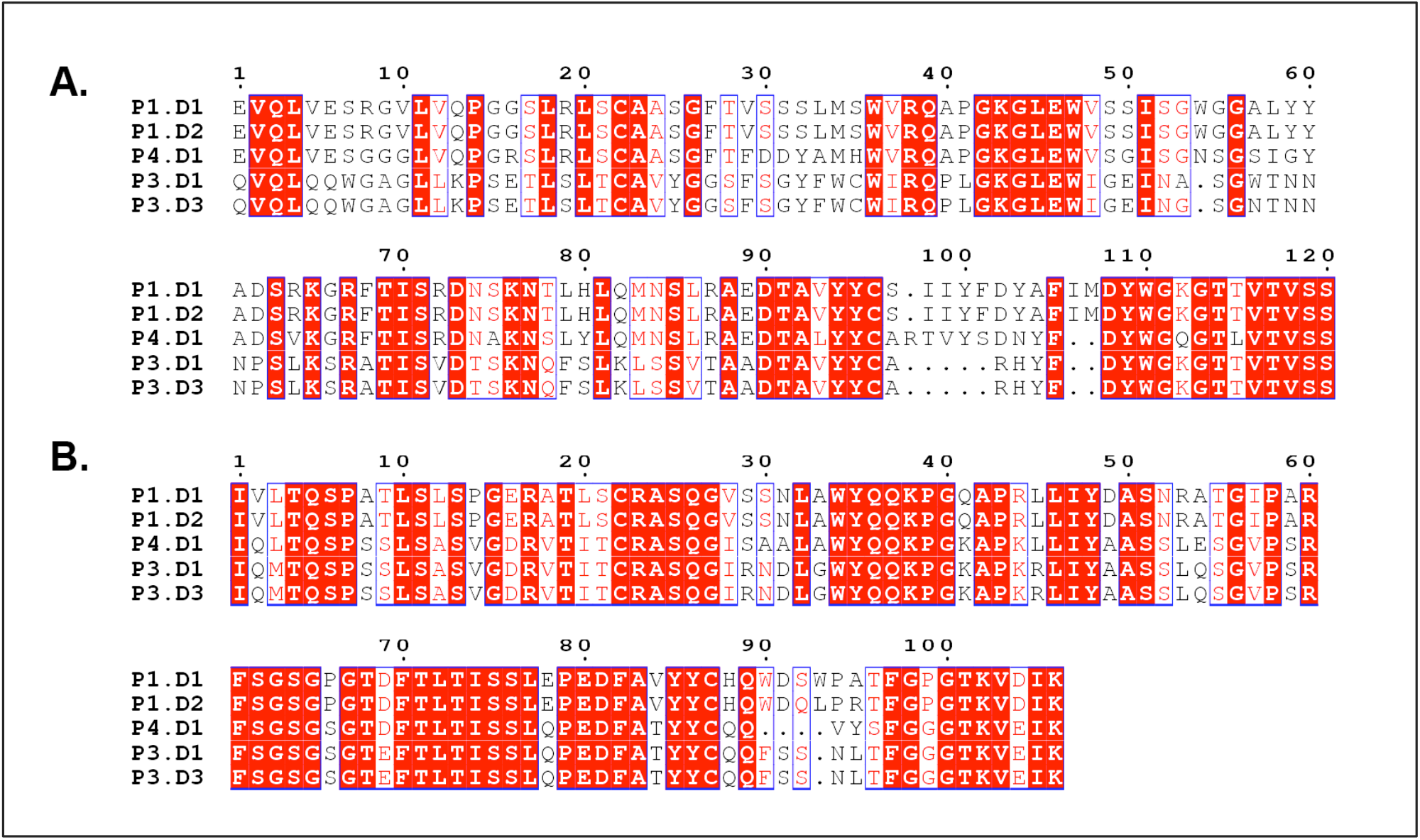
Sequence alignment of Heavy chain sequences (panel A) and light chain sequences (panel B) of the five pareto optimal affinity matured sequences. Conserved positions are outlined in blue boxes. Positions with same residues across designs are highlighted in red and positions with different amino acid but similar residue type are highlighted in white with red text.

Antibodies that strongly bind to the RBD but do not inhibit hACE2 binding have been shown to be neutralizing for SARS-CoV-2 (47D11^14^) and for SARS-CoV-1 (CR3022 in combination with CR3014^32^). The mechanisms of neutralization of such antibodies are not completely known^14^. It is possible that upon binding, these antibodies perturb the interaction network of the RBD with hACE2 thereby rendering RBD-hACE2 binding less effective. In addition, Daniel et al^36^ showed that nanobody VHH-72 raised against SARS-CoV-1 had a neutralizing effect despite binding to an epitope that does not overlap with the hACE2 residue binding domain. By fusing VHH-72 with a human IgG1 they demonstrated SARS-CoV-2 neutralizing activity. They hypothesized that binding of the nanobody with the trimeric spike protein may disrupt conformational dynamics and consequently prevent binding to hACE2.

In comparison, our design P1.D1 forms strong contacts (see Table 2) with many residues of the RBD which in turn also indirectly interact with hACE2 (see Figure 5). For example, residues L455 and T470 of the RBD are in contact with both hACE2 contacting RBD residues Y449, F490 and P1.D1 contacting RBD residues Y351, I465. By perturbing the inter-residue interaction network of RBD-hACE2 it is plausible that a neutralizing effect can be achieved.

**Figure 5.**
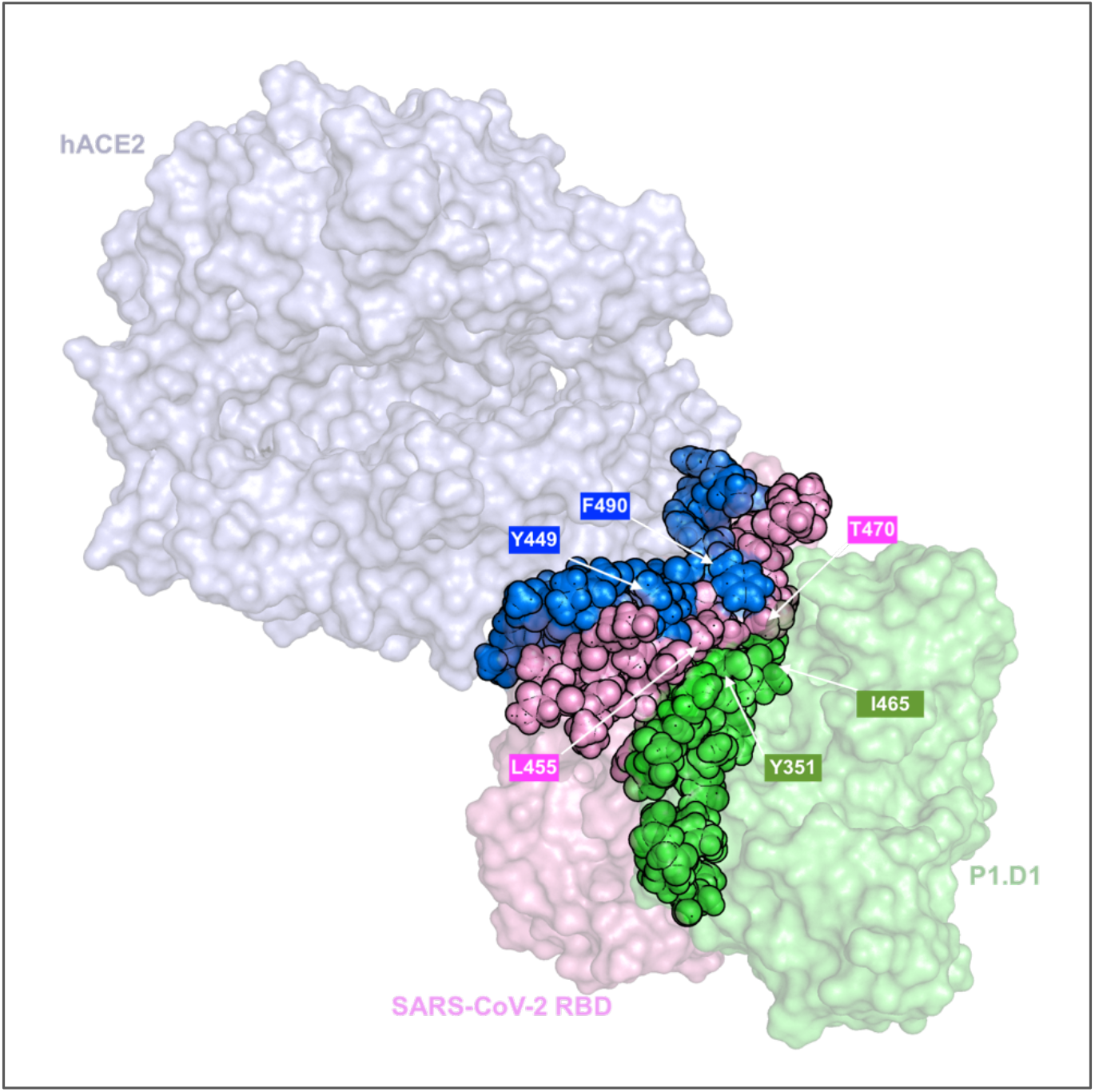
Space filling plot of the P1.D1-RBD complex superimposed on the hACE2-RBD complex. In blue are shown the RBD residues in contact with hACE2 and in green the RBD residues in contact with the P1.D1 designed antibody. In pink are shown all the RBD residues that are in contact with both hACE2 contacting RBD residues and P1.D1 contacting RBD residues. A few residues that are a part of these inter-residue contacts are labelled.

We also carried out an all-atom Molecular Dynamics (MD) simulation of the best binding design P1.D1 in complex with the RBD of the SARS-CoV-2 spike protein to assess the stability of the complex. Preliminary results for a 50ns trajectory counted an average of ~4 hydrogen bonds (st. dev: 1.5) present at the antibody-antigen interface (Supp. Info. S5 for further details). This is quite encouraging, as in an earlier study^37^, MD simulation of the hACE2 receptor in complex with the spike protein RBD reported an average of only 2.7 hydrogen bonds at the interface. This implies that this design has the potential to competitively bind the RBD of the SARS-CoV-2 spike protein potentially sequestering it from hACE2. This is also corroborated by the Rosetta binding energy value of around −48.3 kcal/mol^1^ calculated for the spike protein RBD with hACE2 (from PDB-id: 6lzg) which is weaker by over 7 kcal/mol compared to designs P1.D1 and P1.D2. Finally, it is important to stress that our designs rely on the accuracy of the Rosetta energy function to recapitulate experimental affinities and that carrying out experimental binding assays are needed to confirm or refute these findings.

## Summary

In summary, the goal of this computational analysis was to assess the range of possible antibody designs that can affect binding with the viral spike protein by interacting with residues involved in hACE2 binding. We reported on *de novo* prototype variable regions targeting the most solvent accessible seven-residue epitope in the spike and their (computationally) affinity matured sequences with the lowest Rosetta binding energies. Designs were rank ordered not only in terms of their Rosetta binding energy but also their humanness score metric H-score. We reported complete amino acid sequences for the 106 affinity matured designs as well as the five prototype sequences and V*, CDR3, and J* parts used. Importantly, we would like to note that high affinities of designed antibodies, as modeled using the Rosetta binding energy function, need not necessarily translate to therapeutic effectiveness. The exact mechanisms underlying the therapeutic action of monoclonal antibodies are quite complex and often only partially understood. Nevertheless, we hope that this analysis and data will contribute an important piece to help inform the discovery of high affinity mAb against SARS-CoV-2.

## Methods

### Antibody design in OptMAVEn-2.0

The initial antibody variable domain sequences were predicted using *de novo* antibody design software tool, OptMAVEn-2.0^2^. Using an interatomic clash-cutoff of 1.25 Å, 173 antigen poses were sampled, and each of which yielded a successful (not necessarily unique) antibody design targeted at the seven most solvent accessible hACE2-binding residues of SARS-CoV-2 spike RBD.

Prior to identifying antibody sequences complimentary to the epitopes, OptMAVEn-2.0 first minimizes the z-coordinate of the epitopes, with their collective centroid set at origin, to allow the *de novo* designed antibody regions (see Supp. Info. S1 for link to entire MAPs fragment library) to bind from the bottom. Next, an ensemble of starting antigen poses is generated by a combination of discrete rotations (about the z-axis) and translations (in x, y, and z) – each of which are subsequently passed into the antibody design step. We started out with 3234 such antigen poses for the SARS-CoV-2 spike protein with the epitopes occupying the most negative z-axis coordinates.

### Affinity maturation design in Rosetta

The affinity maturation protocol consisted of an initial refinement of the complex by RosettaDock^38^ followed by three iterations of backbone perturbation using RosettaBackrub^39^, interface design using RosettaDesign^40^ and rotamer repacking of the complex using a Monte Carlo based simulated annealing algorithm^41,42^. During the Rosetta affinity maturation, only amino acids in the variable region within 5 Å from any epitope residue are allowed to mutate. Each affinity matured designed complex was relaxed using FastRelax (with constraints) 10 times and energy minimized (using *Minimize*). For each of these relaxed poses, the binding energy (dG_separated) was calculated using the *InterfaceAnalyzer*^31^ application. The entire protocol was implemented in RosettaScripts^43^ using the REF2015 energy function^30^ (see Supp. info. S6 for further details). This computational protocol was executed for 8,000 affinity matured sequence-design cycles. The top five variable region designs which show strong interaction energy scores with the viral spike and low immunogenicity (high H-scores) were further investigated to glean insight on the biophysics of interactions at the residue level.

## Author contributions

RC designed and performed the OptMAVEn experiments. VSB designed and performed the affinity maturation experiments, MD simulations and related analyses. VSB, RC and CDM wrote the manuscript.

## Acknowledgements

This activity was partially enabled by research conducted within the Center for Bioenergy Innovation of US Department of Energy (DE-SC0018420) and US National Science Foundation (NSF) grant CBET1703274.

All computations were performed using the High-Performance Computing facility at The Inistitute for Computational and Data Sciences Advanced Cyber Infrastructure (ICDS-ACI) at Pennsylvania State University.

## Notes

### Competing Interest Statement

The authors have declared no competing interest.

### Summary of Updates

1. Revised ranking of antibody designs by refining Rosetta binding energy score 2. Standardized implementation of H-score 3. Updated literature information 4. Various text and figure improvements throughout the manuscript 5. Added two supplementary figures

