## Supplementary Material for "*De novo* design of high-affinity antibody variable regions (Fv) against the SARS-CoV-2 spike protein"

**S1. MAPs database[1] full structures download link:**

**http://www.maranasgroup.com/submission/maps.htm**

### S2. P1 to P5 prototype fragment sequences and their respective affinity matured designs

*(Prototype fragment sequences highlighted in bold)*

**>P1**
**EVQLVESRGVLVQPGGSLRLSCAASGFTVSSNEMSWVRQAPGKGLEWVSSISGSGGSTYYADSRKGRFTISRDNSKNTLLQMNSLR**
**AEDTAVYYCSIIYFDYAFIMDYWGKGTTVTVSSEIVLTQSPATLSLSPGERATLSCRASQGVSSNLAWYQQKPGQAPRLLIYDASN**
**RATGIPARFSGSGPGTDFTLTISSLEPEDFAVYYCQWSSYPWTFGPGTKVDIK**

>P1.D1
EVQLVESRGVLVQPGGSLRLSCAASGFTVSSSLMSWVRQAPGKGLEWVSSISGWGGALYYADSRKGRFTISRDNSKNTLHLQMNSL
RAEDTAVYYCSIIYFDYAFIMDYWGKGTTVTVSSIVLTQSPATLSLSPGERATLSCRASQGVSSNLAWYQQKPGQAPRLLIYDASN
RATGIPARFSGSGPGTDFTLTISSLEPEDFAVYYCHQWDSWPATFGPGTKVDIK
>P1.D2
EVQLVESRGVLVQPGGSLRLSCAASGFTVSSSLMSWVRQAPGKGLEWVSSISGWGGALYYADSRKGRFTISRDNSKNTLHLQMNSL
RAEDTAVYYCSIIYFDYAFIMDYWGKGTTVTVSSIVLTQSPATLSLSPGERATLSCRASQGVSSNLAWYQQKPGQAPRLLIYDASN
RATGIPARFSGSGPGTDFTLTISSLEPEDFAVYYCHQWDQLPRTFGPGTKVDIK
>P1.D3
EVQLVESRGVLVQPGGSLRLSCAASGFTVSSSLMSWVRQAPGKGLEWVSSISGWGGATYYADSRKGRFTISRDNSKNTLHLQMNSL
RAEDTAVYYCSIIYFDYAFIMDYWGKGTTVTVSSIVLTQSPATLSLSPGERATLSCRASQGVSSNLAWYQQKPGQAPRLLIYDASN
RATGIPARFSGSGPGTDFTLTISSLEPEDFAVYYCHQWDSIPWTFGPGTKVDIK
>P1.D4
EVQLVESRGVLVQPGGSLRLSCAASGFTVSSSLMSWVRQAPGKGLEWVSSISGWGGALYYADSRKGRFTISRDNSKNTLHLQMNSL
RAEDTAVYYCSIIYFLYAFIMDYWGKGTTVTVSSIVLTQSPATLSLSPGERATLSCRASQGVSSNLAWYQQKPGQAPRLLIYDASN
RATGIPARFSGSGPGTDFTLTISSLEPEDFAVYYCHQWDSIPWTFGPGTKVDIK
>P1.D5
EVQLVESRGVLVQPGGSLRLSCAASGFTVSSSLMSWVRQAPGKGLEWVSSISGWGGALYYADSRKGRFTISRDNSKNTLHLQMNSL
RAEDTAVYYCSIIYFDYAFIMDYWGKGTTVTVSSIVLTQSPATLSLSPGERATLSCRASQGVSSNLAWYQQKPGQAPRLLIYDASN
RATGIPARFSGSGPGTDFTLTISSLEPEDFAVYYCHQWDSYPWTFGPGTKVDIK
>P1.D6
EVQLVESRGVLVQPGGSLRLSCAASGFTVSSSLMSWVRQAPGKGLEWVSSISGWGGALYYADSRKGRFTISRDNSKNTLHLQMNSL
RAEDTAVYYCSIIYFDYAFIMDYWGKGTTVTVSSIVLTQSPATLSLSPGERATLSCRASQGVSSNLAWYQQKPGQAPRLLIYDASN
RATGIPARFSGSGPGTDFTLTISSLEPEDFAVYYCHQWDSIPWTFGPGTKVDIK
>P1.D7
EVQLVESRGVLVQPGGSLRLSCAASGFTVSSSLMSWVRQAPGKGLEWVSSISGWGGALYYADSRKGRFTISRDNSKNTLHLQMNSL
RAEDTAVYYCSIIYFDYAFIMDYWGKGTTVTVSSIVLTQSPATLSLSPGERATLSCRASQGVSSNLAWYQQKPGQAPRLLIYDASN
RATGIPARFSGSGPGTDFTLTISSLEPEDFAVYYCHQWDCWPATFGPGTKVDIK
>P1.D8
EVQLVESRGVLVQPGGSLRLSCAASGFTVSSSLMSWVRQAPGKGLEWVSSISGWGGALYYADSRKGRFTISRDNSKNTLHLQMNSL
RAEDTAVYYCSIIYFDYAFIMDYWGKGTTVTVSSIVLTQSPATLSLSPGERATLSCRASQGVSSNLAWYQQKPGQAPRLLIYDASN
RATGIPARFSGSGPGTDFTLTISSLEPEDFAVYYCHQWDSLPATFGPGTKVDIK
>P1.D9
EVQLVESRGVLVQPGGSLRLSCAASGFTVSSSLMSWVRQAPGKGLEWVSSISGIGGALYYADSRKGRFTISRDNSKNTLHLQMNSL
RAEDTAVYYCSIIYFDYAFIMDYWGKGTTVTVSSIVLTQSPATLSLSPGERATLSCRASQGVSSNLAWYQQKPGQAPRLLIYDASN
RATGIPARFSGSGPGTDFTLTISSLEPEDFAVYYCHQWDSIPWTFGPGTKVDIK
>P1.D10
EVQLVESRGVLVQPGGSLRLSCAASGFTVSSSLMSWVRQAPGKGLEWVSSISGWGGALYYADSRKGRFTISRDNSKNTLHLQMNSL
RAEDTAVYYCSIIYFGYAFIMDYWGKGTTVTVSSIVLTQSPATLSLSPGERATLSCRASQGVSSNLAWYQQKPGQAPRLLIYDASN
RATGIPARFSGSGPGTDFTLTISSLEPEDFAVYYCHQWDAIPWTFGPGTKVDIK
>P1.D11
EVQLVESRGVLVQPGGSLRLSCAASGFTVSSSLMSWVRQAPGKGLEWVSSISGWGGALYYADSRKGRFTISRDNSKNTLHLQMNSL
RAEDTAVYYCSIIYFDYAFIMDYWGKGTTVTVSSIVLTQSPATLSLSPGERATLSCRASQGVSSNLAWYQQKPGQAPRLLIYDASN
RATGIPARFSGSGPGTDFTLTISSLEPEDFAVYYCHQWDSMPWTFGPGTKVDIK
>P1.D12
EVQLVESRGVLVQPGGSLRLSCAASGFTVSSSLMSWVRQAPGKGLEWVSSISGYGGALYYADSRKGRFTISRDNSKNTLHLQMNSL
RAEDTAVYYCSIIYFDYAFIMDYWGKGTTVTVSSIVLTQSPATLSLSPGERATLSCRASQGVSSNLAWYQQKPGQAPRLLIYDASN
RATGIPARFSGSGPGTDFTLTISSLEPEDFAVYYCHQWDSIPWTFGPGTKVDIK
>P1.D13
EVQLVESRGVLVQPGGSLRLSCAASGFTVSSSLMSWVRQAPGKGLEWVSSISGWGGALYYADSRKGRFTISRDNSKNTLHLQMNSL
RAEDTAVYYCSIIYFDYAFIMDYWGKGTTVTVSSIVLTQSPATLSLSPGERATLSCRASQGVSSNLAWYQQKPGQAPRLLIYDASN
RATGIPARFSGSGPGTDFTLTISSLEPEDFAVYYCHQWDAWPATFGPGTKVDIK
>P1.D14
EVQLVESRGVLVQPGGSLRLSCAASGFTVSSGEMSWVRQAPGKGLEWVSSISGWGGALYYADSRKGRFTISRDNSKNTLHLQMNSL
RAEDTAVYYCSIIYFDYAFIMDYWGKGTTVTVSSIVLTQSPATLSLSPGERATLSCRASQGVSSNLAWYQQKPGQAPRLLIYDASN
RATGIPARFSGSGPGTDFTLTISSLEPEDFAVYYCHQWDSLPATFGPGTKVDIK

>P1.D15
EVQLVESRGVLVQPGGSLRLSCAASGFTVSSGNMSWVRQAPGKGLEWVSSISGIGGALYYADSRKGRFTISRDNSKNTLHLQMNSL
RAEDTAVYYCSIIYFDYAFIMDYWGKGTTVTVSSIVLTQSPATLSLSPGERATLSCRASQGVSSNLAWYQQKPGQAPRLLIYDASN
RATGIPARFSGSGPGTDFTLTISSLEPEDFAVYYCHQWDAWPATFGPGTKVDIK
>P1.D16
EVQLVESRGVLVQPGGSLRLSCAASGFTVSSSLMSWVRQAPGKGLEWVSSISGWGGALYYADSRKGRFTISRDNSKNTLHLQMNSL
RAEDTAVYYCSIIYFGYAFIMDYWGKGTTVTVSSIVLTQSPATLSLSPGERATLSCRASQGVSSNLAWYQQKPGQAPRLLIYDASN
RATGIPARFSGSGPGTDFTLTISSLEPEDFAVYYCHQWDSWPATFGPGTKVDIK
>P1.D17
EVQLVESRGVLVQPGGSLRLSCAASGFTVSSSLMSWVRQAPGKGLEWVSSISGWGGALIYADSRKGRFTISRDNSKNTLHLQMNSL
RAEDTAVYYCSIIYFDYAFIMDYWGKGTTVTVSSIVLTQSPATLSLSPGERATLSCRASQGVSSNLAWYQQKPGQAPRLLIYDASN
RATGIPARFSGSGPGTDFTLTISSLEPEDFAVYYCHQWDAWPATFGPGTKVDIK
>P1.D18
EVQLVESRGVLVQPGGSLRLSCAASGFTVSSGEMSWVRQAPGKGLEWVSSISGYGGALYYADSRKGRFTISRDNSKNTLHLQMNSL
RAEDTAVYYCSIIYFDYAFIMDYWGKGTTVTVSSIVLTQSPATLSLSPGERATLSCRASQGVSSNLAWYQQKPGQAPRLLIYDASN
RATGIPARFSGSGPGTDFTLTISSLEPEDFAVYYCHQWDSLPATFGPGTKVDIK
>P1.D19
EVQLVESRGVLVQPGGSLRLSCAASGFTVSSGLMSWVRQAPGKGLEWVSSISGYGGALYYADSRKGRFTISRDNSKNTLHLQMNSL
RAEDTAVYYCSIIYFDYAFIMDYWGKGTTVTVSSIVLTQSPATLSLSPGERATLSCRASQGVSSNLAWYQQKPGQAPRLLIYDASN
RATGIPARFSGSGPGTDFTLTISSLEPEDFAVYYCHQWDSWPATFGPGTKVDIK
>P1.D20
EVQLVESRGVLVQPGGSLRLSCAASGFTVSSSLMSWVRQAPGKGLEWVSSISGWGGALIYADSRKGRFTISRDNSKNTLHLQMNSL
RAEDTAVYYCSIIYFDYAFIMDYWGKGTTVTVSSIVLTQSPATLSLSPGERATLSCRASQGVSSNLAWYQQKPGQAPRLLIYDASN
RATGIPARFSGSGPGTDFTLTISSLEPEDFAVYYCHQWDSYPWTFGPGTKVDIK
>P1.D21
EVQLVESRGVLVQPGGSLRLSCAASGFTVSSGNMSWVRQAPGKGLEWVSSISGIGGALYYADSRKGRFTISRDNSKNTLHLQMNSL
RAEDTAVYYCSIIYFDYAFIMDYWGKGTTVTVSSIVLTQSPATLSLSPGERATLSCRASQGVSSNLAWYQQKPGQAPRLLIYDASN
RATGIPARFSGSGPGTDFTLTISSLEPEDFAVYYCHQWDSWPATFGPGTKVDIK
>P1.D22
EVQLVESRGVLVQPGGSLRLSCAASGFTVSSSLMSWVRQAPGKGLEWVSSISGWGGALIYADSRKGRFTISRDNSKNTLHLQMNSL
RAEDTAVYYCSIIYFDYAFIMDYWGKGTTVTVSSIVLTQSPATLSLSPGERATLSCRASQGVSSNLAWYQQKPGQAPRLLIYDASN
RATGIPARFSGSGPGTDFTLTISSLEPEDFAVYYCHQWDSWPATFGPGTKVDIK
>P1.D23
EVQLVESRGVLVQPGGSLRLSCAASGFTVSSSLMSWVRQAPGKGLEWVSSISGWGGALYYADSRKGRFTISRDNSKNTLHLQMNSL
RAEDTAVYYCSIIYFDYAFIMDYWGKGTTVTVSSIVLTQSPATLSLSPGERATLSCRASQGVSSNLAWYQQKPGQAPRLLIYDASN
RATGIPARFSGSGPGTDFTLTISSLEPEDFAVYYCHQWASIPWTFGPGTKVDIK
>P1.D24
EVQLVESRGVLVQPGGSLRLSCAASGFTVSSGEMSWVRQAPGKGLEWVSSISGSGGALYYADSRKGRFTISRDNSKNTLHLQMNSL
RAEDTAVYYCSIIYFDYAFIMDYWGKGTTVTVSSIVLTQSPATLSLSPGERATLSCRASQGVSSNLAWYQQKPGQAPRLLIYDASN
RATGIPARFSGSGPGTDFTLTISSLEPEDFAVYYCHQWDSLPATFGPGTKVDIK
>P1.D25
EVQLVESRGVLVQPGGSLRLSCAASGFTVSSGEMSWVRQAPGKGLEWVSSISGTGGALYYADSRKGRFTISRDNSKNTLHLQMNSL
RAEDTAVYYCSIIYFDYAFIMDYWGKGTTVTVSSIVLTQSPATLSLSPGERATLSCRASQGVSSNLAWYQQKPGQAPRLLIYDASN
RATGIPARFSGSGPGTDFTLTISSLEPEDFAVYYCHQWDSWPATFGPGTKVDIK
>P1.D26
EVQLVESRGVLVQPGGSLRLSCAASGFTVSSSLMSWVRQAPGKGLEWVSSISGWGGALYYADSRKGRFTISRDNSKNTLHLQMNSL
RAEDTAVYYCSIIYFDYAFIMDYWGKGTTVTVSSIVLTQSPATLSLSPGERATLSCRASQGVSSNLAWYQQKPGQAPRLLIYDASN
RATGIPARFSGSGPGTDFTLTISSLEPEDFAVYYCHQWSSIPWTFGPGTKVDIK
>P1.D27
EVQLVESRGVLVQPGGSLRLSCAASGFTVSSGLMSWVRQAPGKGLEWVSSISGIGGALIYADSRKGRFTISRDNSKNTLHLQMNSL
RAEDTAVYYCSIIYFDYAFIMDYWGKGTTVTVSSIVLTQSPATLSLSPGERATLSCRASQGVSSNLAWYQQKPGQAPRLLIYDASN
RATGIPARFSGSGPGTDFTLTISSLEPEDFAVYYCHQWDSWPATFGPGTKVDIK
>P1.D28
EVQLVESRGVLVQPGGSLRLSCAASGFTVSSSNMSWVRQAPGKGLEWVSSISGYGGALIYADSRKGRFTISRDNSKNTLHLQMNSL
RAEDTAVYYCSIIYFDYAFIMDYWGKGTTVTVSSIVLTQSPATLSLSPGERATLSCRASQGVSSNLAWYQQKPGQAPRLLIYDASN
RATGIPARFSGSGPGTDFTLTISSLEPEDFAVYYCHQWDSWPATFGPGTKVDIK
>P1.D29
EVQLVESRGVLVQPGGSLRLSCAASGFTVSSTEMSWVRQAPGKGLEWVSSISGYGGALIYADSRKGRFTISRDNSKNTLHLQMNSL
RAEDTAVYYCSIIYFDYAFIMDYWGKGTTVTVSSIVLTQSPATLSLSPGERATLSCRASQGVSSNLAWYQQKPGQAPRLLIYDASN
RATGIPARFSGSGPGTDFTLTISSLEPEDFAVYYCHQWDSLPATFGPGTKVDIK
>P1.D30
EVQLVESRGVLVQPGGSLRLSCAASGFTVSSSLMSWVRQAPGKGLEWVSSISGWGGALIYADSRKGRFTISRDNSKNTLHLQMNSL
RAEDTAVYYCSIIYFDYAFIMDYWGKGTTVTVSSIVLTQSPATLSLSPGERATLSCRASQGVSSNLAWYQQKPGQAPRLLIYDASN
RATGIPARFSGSGPGTDFTLTISSLEPEDFAVYYCHQWASWPATFGPGTKVDIK

**>P2**
**VQLVESGGGVVQPGRSLRLSCAASGFTFSSYAMWVRQAPGKGLEWVAVISYDGSNKYYADSVKGRFAISRDNSKNTLYLQMNSLRA**
**EDTAVYYCATDDPDAFDIWGQGTLVTVSSEIVLTQSPATLSLSPGERATLSCRASQGVSSNLAWYQQKPGQAPRLLIYDASNRATG**
**IPARFSGSGPGTDFTLTISSLEPEDFAVYYCQQANSFLTFGPGTKVDIK**

>P2.D1
VQLVESGGGVVQPGRSLRLSCAASGFTFSSYAMHWVRQAPGKGLEWVAVIAYDNGYSYYADSVKGRFAISRDNSKNTLYLQMNSLR
AEDTAVYYCATDDPDAFDIWGQGTLVTVSSIVLTQSPATLSLSPGERATLSCRASQGVSSNLAWYQQKPGQAPRLLIYDASNRATG
IPARFSGSGPGTDFTLTISSLEPEDFAVYYCQQAGSFLTFGPGTKVDIK
>P2.D2
VQLVESGGGVVQPGRSLRLSCAASGFTFSSYAMHWVRQAPGKGLEWVAVIAYDNGYAYYADSVKGRFAISRDNSKNTLYLQMNSLR
AEDTAVYYCATDDPDAFDIWGQGTLVTVSSIVLTQSPATLSLSPGERATLSCRASQGVSSNLAWYQQKPGQAPRLLIYDASNRATG
IPARFSGSGPGTDFTLTISSLEPEDFAVYYCQQAGSFLTFGPGTKVDIK
>P2.D3
VQLVESGGGVVQPGRSLRLSCAASGFTFSSYAMHWVRQAPGKGLEWVAVIAYDNGDSYYADSVKGRFAISRDNSKNTLYLQMNSLR
AEDTAVYYCATDDPDAFDIWGQGTLVTVSSIVLTQSPATLSLSPGERATLSCRASQGVSSNLAWYQQKPGQAPRLLIYDASNRATG
IPARFSGSGPGTDFTLTISSLEPEDFAVYYCQQAGSFLTFGPGTKVDIK
>P2.D4
VQLVESGGGVVQPGRSLRLSCAASGFTFSSYAMHWVRQAPGKGLEWVAVIADDNGYAYYADSVKGRFAISRDNSKNTLYLQMNSLR
AEDTAVYYCATDDPDAFDIWGQGTLVTVSSIVLTQSPATLSLSPGERATLSCRASQGVSSNLAWYQQKPGQAPRLLIYDASNRATG
IPARFSGSGPGTDFTLTISSLEPEDFAVYYCQQAGSFLTFGPGTKVDIK
>P2.D5
VQLVESGGGVVQPGRSLRLSCAASGFTFSSYAMHWVRQAPGKGLEWVAVIAADNGYAYYADSVKGRFAISRDNSKNTLYLQMNSLR
AEDTAVYYCATDDPDAFDIWGQGTLVTVSSIVLTQSPATLSLSPGERATLSCRASQGVSSNLAWYQQKPGQAPRLLIYDASNRATG
IPARFSGSGPGTDFTLTISSLEPEDFAVYYCQQAGSFLTFGPGTKVDIK
>P2.D6
VQLVESGGGVVQPGRSLRLSCAASGFTFSSYAMHWVRQAPGKGLEWVAVITGDKGYSYYADSVKGRFAISRDNSKNTLYLQMNSLR
AEDTAVYYCATDDPDAFDIWGQGTLVTVSSIVLTQSPATLSLSPGERATLSCRASQGVSSNLAWYQQKPGQAPRLLIYDASNRATG
IPARFSGSGPGTDFTLTISSLEPEDFAVYYCQQAGSFLTFGPGTKVDIK
>P2.D7
VQLVESGGGVVQPGRSLRLSCAASGFTFSSYAMHWVRQAPGKGLEWVAVIAYDNGYVYYADSVKGRFAISRDNSKNTLYLQMNSLR
AEDTAVYYCATDDPDAFDIWGQGTLVTVSSIVLTQSPATLSLSPGERATLSCRASQGVSSNLAWYQQKPGQAPRLLIYDASNRATG
IPARFSGSGPGTDFTLTISSLEPEDFAVYYCQQAGSFLTFGPGTKVDIK
>P2.D8
VQLVESGGGVVQPGRSLRLSCAASGFTFSSYAMHWVRQAPGKGLEWVAVITGDNGYSYYADSVKGRFAISRDNSKNTLYLQMNSLR
AEDTAVYYCATDDPDAFDIWGQGTLVTVSSIVLTQSPATLSLSPGERATLSCRASQGVSSNLAWYQQKPGQAPRLLIYDASNRATG
IPARFSGSGPGTDFTLTISSLEPEDFAVYYCQQAGSFLTFGPGTKVDIK
>P2.D9
VQLVESGGGVVQPGRSLRLSCAASGFTFSSYAMHWVRQAPGKGLEWVAVIAYDNSYAYYADSVKGRFAISRDNSKNTLYLQMNSLR
AEDTAVYYCATDDPDAFDIWGQGTLVTVSSIVLTQSPATLSLSPGERATLSCRASQGVSSNLAWYQQKPGQAPRLLIYDASNRATG
IPARFSGSGPGTDFTLTISSLEPEDFAVYYCQQAGTFLTFGPGTKVDIK
>P2.D10
VQLVESGGGVVQPGRSLRLSCAASGFTFSSYAMHWVRQAPGKGLEWVAVIAADNGYSYYADSVKGRFAISRDNSKNTLYLQMNSLR
AEDTAVYYCATDDPDAFDIWGQGTLVTVSSIVLTQSPATLSLSPGERATLSCRASQGVSSNLAWYQQKPGQAPRLLIYDASNRATG
IPARFSGSGPGTDFTLTISSLEPEDFAVYYCQQAGSALTFGPGTKVDIK
>P2.D11
VQLVESGGGVVQPGRSLRLSCAASGFTFSSYAMHWVRQAPGKGLEWVAVIAYDNSYSYYADSVKGRFAISRDNSKNTLYLQMNSLR
AEDTAVYYCATDDPDAFDIWGQGTLVTVSSIVLTQSPATLSLSPGERATLSCRASQGVSSNLAWYQQKPGQAPRLLIYDASNRATG
IPARFSGSGPGTDFTLTISSLEPEDFAVYYCQQAGSFLTFGPGTKVDIK
>P2.D12
VQLVESGGGVVQPGRSLRLSCAASGFTFSSYAMHWVRQAPGKGLEWVAVIAYDNSYAYYADSVKGRFAISRDNSKNTLYLQMNSLR
AEDTAVYYCATDDPDAFDIWGQGTLVTVSSIVLTQSPATLSLSPGERATLSCRASQGVSSNLAWYQQKPGQAPRLLIYDASNRATG
IPARFSGSGPGTDFTLTISSLEPEDFAVYYCQQAGSFLTFGPGTKVDIK
>P2.D13
VQLVESGGGVVQPGRSLRLSCAASGFTFSSYAMHWVRQAPGKGLEWVAVIGYDNSYAYYADSVKGRFAISRDNSKNTLYLQMNSLR
AEDTAVYYCATDDPDAFDIWGQGTLVTVSSIVLTQSPATLSLSPGERATLSCRASQGVSSNLAWYQQKPGQAPRLLIYDASNRATG
IPARFSGSGPGTDFTLTISSLEPEDFAVYYCQQAGSFLTFGPGTKVDIK

**>P3**
**QVQLQQWGAGLLKPSETLSLTCAVYGGSFSGYYWCWIRQPLGKGLEWIGEINSGSTNNNPSLKSRATISVDTSKNQFSLKLSSVTA**
**ADTAVYYCARYFDYWGKGTTVTVSSDIQMTQSPSSLSASVGDRVTITCRASQGIRNDLGWYQQKPGKAPKRLIYAASSLQSGVPSR**
**FSGSGSGTEFTLTISSLQPEDFATYYCQQWSSNLTFGGGTKVEIK**

>P3.D1
QVQLQQWGAGLLKPSETLSLTCAVYGGSFSGYFWCWIRQPLGKGLEWIGEINASGWTNNNPSLKSRATISVDTSKNQFSLKLSSVT
AADTAVYYCARHYFDYWGKGTTVTVSSIQMTQSPSSLSASVGDRVTITCRASQGIRNDLGWYQQKPGKAPKRLIYAASSLQSGVPS
RFSGSGSGTEFTLTISSLQPEDFATYYCQQFSSNLTFGGGTKVEIK
>P3.D2
QVQLQQWGAGLLKPSETLSLTCAVYGGSFSGYFWCWIRQPLGKGLEWIGEINHSGNTNNNPSLKSRATISVDTSKNQFSLKLSSVT
AADTAVYYCARHYFDYWGKGTTVTVSSIQMTQSPSSLSASVGDRVTITCRASQGIRNDLGWYQQKPGKAPKRLIYAASSLQSGVPS
RFSGSGSGTEFTLTISSLQPEDFATYYCQCFTSNLTFGGGTKVEIK
>P3.D3
QVQLQQWGAGLLKPSETLSLTCAVYGGSFSGYFWCWIRQPLGKGLEWIGEINGSGNTNNNPSLKSRATISVDTSKNQFSLKLSSVT
AADTAVYYCARHYFDYWGKGTTVTVSSIQMTQSPSSLSASVGDRVTITCRASQGIRNDLGWYQQKPGKAPKRLIYAASSLQSGVPS
RFSGSGSGTEFTLTISSLQPEDFATYYCQQFSSNLTFGGGTKVEIK
>P3.D4
QVQLQQWGAGLLKPSETLSLTCAVYGGSFSGYFWCWIRQPLGKGLEWIGEINASGMTNNNPSLKSRATISVDTSKNQFSLKLSSVT
AADTAVYYCARHYFDYWGKGTTVTVSSIQMTQSPSSLSASVGDRVTITCRASQGIRNDLGWYQQKPGKAPKRLIYAASSLQSGVPS
RFSGSGSGTEFTLTISSLQPEDFATYYCQSSTSNLTFGGGTKVEIK
>P3.D5
QVQLQQWGAGLLKPSETLSLTCAVYGGSFSGYFWCWIRQPLGKGLEWIGEINASGNTNNNPSLKSRATISVDTSKNQFSLKLSSVT
AADTAVYYCARHYFDYWGKGTTVTVSSIQMTQSPSSLSASVGDRVTITCRASQGIRNDLGWYQQKPGKAPKRLIYAASSLQSGVPS
RFSGSGSGTEFTLTISSLQPEDFATYYCQQFSSNLTFGGGTKVEIK
>P3.D6
QVQLQQWGAGLLKPSETLSLTCAVYGGSFSGYFWCWIRQPLGKGLEWIGEINHSGWTNNNPSLKSRATISVDTSKNQFSLKLSSVT
AADTAVYYCARHYFDYWGKGTTVTVSSIQMTQSPSSLSASVGDRVTITCRASQGIRNDLGWYQQKPGKAPKRLIYAASSLQSGVPS
RFSGSGSGTEFTLTISSLQPEDFATYYCQQFSSNLTFGGGTKVEIK
>P3.D7
QVQLQQWGAGLLKPSETLSLTCAVYGGSFSGYFWCWIRQPLGKGLEWIGEINHSGNTNNNPSLKSRATISVDTSKNQFSLKLSSVT
AADTAVYYCARHYFDYWGKGTTVTVSSIQMTQSPSSLSASVGDRVTITCRASQGIRNDLGWYQQKPGKAPKRLIYAASSLQSGVPS
RFSGSGSGTEFTLTISSLQPEDFATYYCQQFSSNLTFGGGTKVEIK
>P3.D8
QVQLQQWGAGLLKPSETLSLTCAVYGGSFSGYFWCWIRQPLGKGLEWIGEINHSGNTNNNPSLKSRATISVDTSKNQFSLKLSSVT
AADTAVYYCARHYFDYWGKGTTVTVSSIQMTQSPSSLSASVGDRVTITCRASQGIRNTLGWYQQKPGKAPKRLIYAASSLQSGVPS
RFSGSGSGTEFTLTISSLQPEDFATYYCQQFSSNLTFGGGTKVEIK
>P3.D9
QVQLQQWGAGLLKPSETLSLTCAVYGGSFSGYFWCWIRQPLGKGLEWIGEINHSGNTNNNPSLKSRATISVDTSKNQFSLKLSSVT
AADTAVYYCARHYFDYWGKGTTVTVSSIQMTQSPSSLSASVGDRVTITCRASQGIRNSLGWYQQKPGKAPKRLIYAASSLQSGVPS
RFSGSGSGTEFTLTISSLQPEDFATYYCQQFSSNLTFGGGTKVEIK
>P3.D10
QVQLQQWGAGLLKPSETLSLTCAVYGGSFSGYFWCWIRQPLGKGLEWIGEINHSGNTNNNPSLKSRATISVDTSKNQFSLKLSSVT
AADTAVYYCARGYFDYWGKGTTVTVSSIQMTQSPSSLSASVGDRVTITCRASQGIRNDLGWYQQKPGKAPKRLIYAASSLQSGVPS
RFSGSGSGTEFTLTISSLQPEDFATYYCQQFSSNLTFGGGTKVEIK
>P3.D11
QVQLQQWGAGLLKPSETLSLTCAVYGGSFSGYFWCWIRQPLGKGLEWIGEINHSGLTNNNPSLKSRATISVDTSKNQFSLKLSSVT
AADTAVYYCARHYFDYWGKGTTVTVSSIQMTQSPSSLSASVGDRVTITCRASQGIRNDLGWYQQKPGKAPKRLIYAASSLQSGVPS
RFSGSGSGTEFTLTISSLQPEDFATYYCQCHSSNLTFGGGTKVEIK

**>P4.D1**
**EVQLVESGGGLVQPGRSLRLSCAASGFTFDDYAMHWVRQAPGKGLEWVSGISSNSGSIGYADSVKGRFTISRDNAKNSLYLQMNSL**
**RAEDTALYYCARTLYSDNYFDYWGQGTLVTVSSIQLTQSPSSLSASVGDRVTITCRASQGISAALAWYQQKPGKAPKLLIYAASSL**
**DSGVPSRFSGSGSGTDFTLTISSLQPEDFATYYCQQVYSFGGGTKVEIK**

>P4.D1
EVQLVESGGGLVQPGRSLRLSCAASGFTFDDYAMHWVRQAPGKGLEWVSGISGNSGSIGYADSVKGRFTISRDNAKNSLYLQMNSL
RAEDTALYYCARTVYSDNYFDYWGQGTLVTVSSIQLTQSPSSLSASVGDRVTITCRASQGISAALAWYQQKPGKAPKLLIYAASSL
ESGVPSRFSGSGSGTDFTLTISSLQPEDFATYYCQQVYSFGGGTKVEIK
>P4.D2
EVQLVESGGGLVQPGRSLRLSCAASGFTFDDYAMHWVRQAPGKGLEWVSGISGNSGSIGYADSVKGRFTISRDNAKNSLYLQMNSL
RAEDTALYYCARTLYSDNYFDYWGQGTLVTVSSIQLTQSPSSLSASVGDRVTITCRASQGISAALAWYQQKPGKAPKLLIYAASSL
ASGVPSRFSGSGSGTDFTLTISSLQPEDFATYYCQQVYSFGGGTKVEIK
>P4.D3

EVQLVESGGGLVQPGRSLRLSCAASGFTFDDYAMHWVRQAPGKGLEWVSGISSNSGSIGYADSVKGRFTISRDNAKNSLYLQMNSL
RAEDTALYYCARTLDKDNYFDYWGQGTLVTVSSIQLTQSPSSLSASVGDRVTITCRASQGISAALAWYQQKPGKAPKLLIYAASSL
DSGVPSRFSGSGSGTDFTLTISSLQPEDFATYYCQQVYSFGGGTKVEIK
>P4.D4
EVQLVESGGGLVQPGRSLRLSCAASGFTFDDYAMHWVRQAPGKGLEWVSGISSNSGSIGYADSVKGRFTISRDNAKNSLYLQMNSL
RAEDTALYYCARTVAGDNYFDYWGQGTLVTVSSIQLTQSPSSLSASVGDRVTITCRASQGISAALAWYQQKPGKAPKLLIYDASSL
ESGVPSRFSGSGSGTDFTLTISSLQPEDFATYYCQQVYSFGGGTKVEIK
>P4.D5
EVQLVESGGGLVQPGRSLRLSCAASGFTFDDYAMHWVRQAPGKGLEWVSGISSNSGSIGYADSVKGRFTISRDNAKNSLYLQMNSL
RAEDTALYYCARTQDGDNYFDYWGQGTLVTVSSIQLTQSPSSLSASVGDRVTITCRASQGISAALAWYQQKPGKAPKLLIYAASSL
ESGVPSRFSGSGSGTDFTLTISSLQPEDFATYYCQQVYSFGGGTKVEIK
>P4.D6
EVQLVESGGGLVQPGRSLRLSCAASGFTFDDYAMHWVRQAPGKGLEWVSGISSNSGSIGYADSVKGRFTISRDNAKNSLYLQMNSL
RAEDTALYYCARTVDKSNYFDYWGQGTLVTVSSIQLTQSPSSLSASVGDRVTITCRASQGISAALAWYQQKPGKAPKLLIYDASSL
ESGVPSRFSGSGSGTDFTLTISSLQPEDFATYYCQQVYSFGGGTKVEIK
>P4.D7
EVQLVESGGGLVQPGRSLRLSCAASGFTFDDYAMHWVRQAPGKGLEWVSGISSNSGSIGYADSVKGRFTISRDNAKNSLYLQMNSL
RAEDTALYYCARTVNKSNYFDYWGQGTLVTVSSIQLTQSPSSLSASVGDRVTITCRASQGISAALAWYQQKPGKAPKLLIYDASSL
DSGVPSRFSGSGSGTDFTLTISSLQPEDFATYYCQQVYSFGGGTKVEIK
>P4.D8
EVQLVESGGGLVQPGRSLRLSCAASGFTFDDYAMHWVRQAPGKGLEWVSGISSNSGSIGYADSVKGRFTISRDNAKNSLYLQMNSL
RAEDTALYYCARTIDKSNYFDYWGQGTLVTVSSIQLTQSPSSLSASVGDRVTITCRASQGISAALAWYQQKPGKAPKLLIYDASSL
ESGVPSRFSGSGSGTDFTLTISSLQPEDFATYYCQQVYSFGGGTKVEIK
>P4.D9
EVQLVESGGGLVQPGRSLRLSCAASGFTFDDYAMHWVRQAPGKGLEWVSGISSNSGSIGYADSVKGRFTISRDNAKNSLYLQMNSL
RAEDTALYYCARTLDNANYFDYWGQGTLVTVSSIQLTQSPSSLSASVGDRVTITCRASQGISAALAWYQQKPGKAPKLLIYAASSL
DSGVPSRFSGSGSGTDFTLTISSLQPEDFATYYCQQVYSFGGGTKVEIK
>P4.D10
EVQLVESGGGLVQPGRSLRLSCAASGFTFDDYAMHWVRQAPGKGLEWVSGISSNSGSIGYADSVKGRFTISRDNAKNSLYLQMNSL
RAEDTALYYCARTLDKDNYFDYWGQGTLVTVSSIQLTQSPSSLSASVGDRVTITCRASQGISAALAWYQQKPGKAPKLLIYAASSL
TSGVPSRFSGSGSGTDFTLTISSLQPEDFATYYCQQVYSFGGGTKVEIK
>P4.D11
EVQLVESGGGLVQPGRSLRLSCAASGFTFDDYAMHWVRQAPGKGLEWVSGISSNSGSIGYADSVKGRFTISRDNAKNSLYLQMNSL
RAEDTALYYCARTLDKSNYFDYWGQGTLVTVSSIQLTQSPSSLSASVGDRVTITCRASQGISAALAWYQQKPGKAPKLLIYDASSL
DSGVPSRFSGSGSGTDFTLTISSLQPEDFATYYCQQVYSFGGGTKVEIK
>P4.D12
EVQLVESGGGLVQPGRSLRLSCAASGFTFDDYAMHWVRQAPGKGLEWVSGISSNSGSIGYADSVKGRFTISRDNAKNSLYLQMNSL
RAEDTALYYCARTVDKANYFDYWGQGTLVTVSSIQLTQSPSSLSASVGDRVTITCRASQGISAALAWYQQKPGKAPKLLIYDASSL
TSGVPSRFSGSGSGTDFTLTISSLQPEDFATYYCQQVYSFGGGTKVEIK
>P4.D13
EVQLVESGGGLVQPGRSLRLSCAASGFTFDDYAMHWVRQAPGKGLEWVSGISSNSGSIGYADSVKGRFTISRDNAKNSLYLQMNSL
RAEDTALYYCARTLSNTNYFDYWGQGTLVTVSSIQLTQSPSSLSASVGDRVTITCRASQGISAALAWYQQKPGKAPKLLIYDASSL
ESGVPSRFSGSGSGTDFTLTISSLQPEDFATYYCQQVYSFGGGTKVEIK
>P4.D14
EVQLVESGGGLVQPGRSLRLSCAASGFTFDDYAMHWVRQAPGKGLEWVSGISSNSGSIGYADSVKGRFTISRDNAKNSLYLQMNSL
RAEDTALYYCARTVDGANYFDYWGQGTLVTVSSIQLTQSPSSLSASVGDRVTITCRASQGISAALAWYQQKPGKAPKLLIYAASSL
HSGVPSRFSGSGSGTDFTLTISSLQPEDFATYYCQQVYSFGGGTKVEIK
>P4.D15
EVQLVESGGGLVQPGRSLRLSCAASGFTFDDYAMHWVRQAPGKGLEWVSGISSNSGSIGYADSVKGRFTISRDNAKNSLYLQMNSL
RAEDTALYYCARTLDRSNYFDYWGQGTLVTVSSIQLTQSPSSLSASVGDRVTITCRASQGISAALAWYQQKPGKAPKLLIYDASSL
ESGVPSRFSGSGSGTDFTLTISSLQPEDFATYYCQQVYSFGGGTKVEIK
>P4.D16
EVQLVESGGGLVQPGRSLRLSCAASGFTFDDYAMHWVRQAPGKGLEWVSGISSNSGSIGYADSVKGRFTISRDNAKNSLYLQMNSL
RAEDTALYYCARTIDKANYFDYWGQGTLVTVSSIQLTQSPSSLSASVGDRVTITCRASQGISAALAWYQQKPGKAPKLLIYAASSL
ASGVPSRFSGSGSGTDFTLTISSLQPEDFATYYCQQVYSFGGGTKVEIK
>P4.D17
EVQLVESGGGLVQPGRSLRLSCAASGFTFDDYAMHWVRQAPGKGLEWVSGISSNSGSIGYADSVKGRFTISRDNAKNSLYLQMNSL
RAEDTALYYCARTVDKANYFDYWGQGTLVTVSSIQLTQSPSSLSASVGDRVTITCRASQGISAALAWYQQKPGKAPKLLIYAASSL
DSGVPSRFSGSGSGTDFTLTISSLQPEDFATYYCQQVYSFGGGTKVEIK
>P4.D18
EVQLVESGGGLVQPGRSLRLSCAASGFTFDDYAMHWVRQAPGKGLEWVSGISSNSGSIGYADSVKGRFTISRDNAKNSLYLQMNSL
RAEDTALYYCARTIDGSNYFDYWGQGTLVTVSSIQLTQSPSSLSASVGDRVTITCRASQGISAALAWYQQKPGKAPKLLIYDASSL
ESGVPSRFSGSGSGTDFTLTISSLQPEDFATYYCQQVYSFGGGTKVEIK
>P4.D19

EVQLVESGGGLVQPGRSLRLSCAASGFTFDDYAMHWVRQAPGKGLEWVSGISSNSGSIGYADSVKGRFTISRDNAKNSLYLQMNSL
RAEDTALYYCARTLDGANYFDYWGQGTLVTVSSIQLTQSPSSLSASVGDRVTITCRASQGISAALAWYQQKPGKAPKLLIYDASSL
LSGVPSRFSGSGSGTDFTLTISSLQPEDFATYYCQQVYSFGGGTKVEIK
>P4.D20
EVQLVESGGGLVQPGRSLRLSCAASGFTFDDYAMHWVRQAPGKGLEWVSGISSNSGSIGYADSVKGRFTISRDNAKNSLYLQMNSL
RAEDTALYYCARTVDKSNYFDYWGQGTLVTVSSIQLTQSPSSLSASVGDRVTITCRASQGISAALAWYQQKPGKAPKLLIYDASSL
DSGVPSRFSGSGSGTDFTLTISSLQPEDFATYYCQQVYSFGGGTKVEIK
>P4.D21
EVQLVESGGGLVQPGRSLRLSCAASGFTFDDYAMHWVRQAPGKGLEWVSGISGNSGSIGYADSVKGRFTISRDNAKNSLYLQMNSL
RAEDTALYYCARTVNKANYFDYWGQGTLVTVSSIQLTQSPSSLSASVGDRVTITCRASQGISAALAWYQQKPGKAPKLLIYAASSL
TSGVPSRFSGSGSGTDFTLTISSLQPEDFATYYCQQVYSFGGGTKVEIK
>P4.D22
EVQLVESGGGLVQPGRSLRLSCAASGFTFDDYAMHWVRQAPGKGLEWVSGISSNSGSIGYADSVKGRFTISRDNAKNSLYLQMNSL
RAEDTALYYCARTLDGANYFDYWGQGTLVTVSSIQLTQSPSSLSASVGDRVTITCRASQGISAALAWYQQKPGKAPKLLIYAASSL
DSGVPSRFSGSGSGTDFTLTISSLQPEDFATYYCQQVYSFGGGTKVEIK
>P4.D23
EVQLVESGGGLVQPGRSLRLSCAASGFTFDDYAMHWVRQAPGKGLEWVSGISSNSGSIGYADSVKGRFTISRDNAKNSLYLQMNSL
RAEDTALYYCARTLDGANYFDYWGQGTLVTVSSIQLTQSPSSLSASVGDRVTITCRASQGISAALAWYQQKPGKAPKLLIYAASSL
ESGVPSRFSGSGSGTDFTLTISSLQPEDFATYYCQQVYSFGGGTKVEIK
>P4.D24
EVQLVESGGGLVQPGRSLRLSCAASGFTFDDYAMHWVRQAPGKGLEWVSGISSNSGSIGYADSVKGRFTISRDNAKNSLYLQMNSL
RAEDTALYYCARTDDKANYFDYWGQGTLVTVSSIQLTQSPSSLSASVGDRVTITCRASQGISAALAWYQQKPGKAPKLLIYDASSL
TSGVPSRFSGSGSGTDFTLTISSLQPEDFATYYCQQVYSFGGGTKVEIK
>P4.D25
EVQLVESGGGLVQPGRSLRLSCAASGFTFDDYAMHWVRQAPGKGLEWVSGISGNSGSIGYADSVKGRFTISRDNAKNSLYLQMNSL
RAEDTALYYCARTLDKANYFDYWGQGTLVTVSSIQLTQSPSSLSASVGDRVTITCRASQGISAALAWYQQKPGKAPKLLIYAASSL
ESGVPSRFSGSGSGTDFTLTISSLQPEDFATYYCQQVYSFGGGTKVEIK
>P4.D26
EVQLVESGGGLVQPGRSLRLSCAASGFTFDDYAMHWVRQAPGKGLEWVSGISGNSGSIGYADSVKGRFTISRDNAKNSLYLQMNSL
RAEDTALYYCARTIDKANYFDYWGQGTLVTVSSIQLTQSPSSLSASVGDRVTITCRASQGISAALAWYQQKPGKAPKLLIYAASSL
TSGVPSRFSGSGSGTDFTLTISSLQPEDFATYYCQQVYSFGGGTKVEIK
>P4.D27
EVQLVESGGGLVQPGRSLRLSCAASGFTFDDYAMHWVRQAPGKGLEWVSGISSNSGSIGYADSVKGRFTISRDNAKNSLYLQMNSL
RAEDTALYYCARTIDKANYFDYWGQGTLVTVSSIQLTQSPSSLSASVGDRVTITCRASQGISAALAWYQQKPGKAPKLLIYAASSL
TSGVPSRFSGSGSGTDFTLTISSLQPEDFATYYCQQVYSFGGGTKVEIK
>P4.D28
EVQLVESGGGLVQPGRSLRLSCAASGFTFDDYAMHWVRQAPGKGLEWVSGISSNSGSIGYADSVKGRFTISRDNAKNSLYLQMNSL
RAEDTALYYCARTLSKANYFDYWGQGTLVTVSSIQLTQSPSSLSASVGDRVTITCRASQGISAALAWYQQKPGKAPKLLIYAASSL
ESGVPSRFSGSGSGTDFTLTISSLQPEDFATYYCQQVYSFGGGTKVEIK
>P4.D29
EVQLVESGGGLVQPGRSLRLSCAASGFTFDDYAMHWVRQAPGKGLEWVSGISSNSGSIGYADSVKGRFTISRDNAKNSLYLQMNSL
RAEDTALYYCARTIDKANYFDYWGQGTLVTVSSIQLTQSPSSLSASVGDRVTITCRASQGISAALAWYQQKPGKAPKLLIYAASSL
DSGVPSRFSGSGSGTDFTLTISSLQPEDFATYYCQQVYSFGGGTKVEIK
>P4.D30
EVQLVESGGGLVQPGRSLRLSCAASGFTFDDYAMHWVRQAPGKGLEWVSGISGNSGSIGYADSVKGRFTISRDNAKNSLYLQMNSL
RAEDTALYYCARTLDKANYFDYWGQGTLVTVSSIQLTQSPSSLSASVGDRVTITCRASQGISAALAWYQQKPGKAPKLLIYAASSL
DSGVPSRFSGSGSGTDFTLTISSLQPEDFATYYCQQVYSFGGGTKVEIK
>P4.D31
EVQLVESGGGLVQPGRSLRLSCAASGFTFDDYAMHWVRQAPGKGLEWVSGISSNSGSIGYADSVKGRFTISRDNAKNSLYLQMNSL
RAEDTALYYCARTVDSANYFDYWGQGTLVTVSSIQLTQSPSSLSASVGDRVTITCRASQGISAALAWYQQKPGKAPKLLIYAASSL
DSGVPSRFSGSGSGTDFTLTISSLQPEDFATYYCQQVYSFGGGTKVEIK
>P4.D32
EVQLVESGGGLVQPGRSLRLSCAASGFTFDDYAMHWVRQAPGKGLEWVSGISSNSGSIGYADSVKGRFTISRDNAKNSLYLQMNSL
RAEDTALYYCARTVDKTNYFDYWGQGTLVTVSSIQLTQSPSSLSASVGDRVTITCRASQGISAALAWYQQKPGKAPKLLIYDASSL
DSGVPSRFSGSGSGTDFTLTISSLQPEDFATYYCQQVYSFGGGTKVEIK
>P4.D33
EVQLVESGGGLVQPGRSLRLSCAASGFTFDDYAMHWVRQAPGKGLEWVSGISSNSGSIGYADSVKGRFTISRDNAKNSLYLQMNSL
RAEDTALYYCARTLDKANYFDYWGQGTLVTVSSIQLTQSPSSLSASVGDRVTITCRASQGISCALAWYQQKPGKAPKLLIYDASSL
TSGVPSRFSGSGSGTDFTLTISSLQPEDFATYYCQQVYSFGGGTKVEIK
>P4.D34
EVQLVESGGGLVQPGRSLRLSCAASGFTFDDYAMHWVRQAPGKGLEWVSGISANSGSIGYADSVKGRFTISRDNAKNSLYLQMNSL
RAEDTALYYCARTLDKANYFDYWGQGTLVTVSSIQLTQSPSSLSASVGDRVTITCRASQGISAALAWYQQKPGKAPKLLIYAASSL
DSGVPSRFSGSGSGTDFTLTISSLQPEDFATYYCQQVYSFGGGTKVEIK
>P4.D35

EVQLVESGGGLVQPGRSLRLSCAASGFTFDDYAMHWVRQAPGKGLEWVSGISSNSGSIGYADSVKGRFTISRDNAKNSLYLQMNSL
RAEDTALYYCARTVDGANYFDYWGQGTLVTVSSIQLTQSPSSLSASVGDRVTITCRASQGISAALAWYQQKPGKAPKLLIYAASSL
ESGVPSRFSGSGSGTDFTLTISSLQPEDFATYYCQQVYSFGGGTKVEIK
>P4.D36
EVQLVESGGGLVQPGRSLRLSCAASGFTFDDYAMHWVRQAPGKGLEWVSGISSNSGSIGYADSVKGRFTISRDNAKNSLYLQMNSL
RAEDTALYYCARTVNNTNYFDYWGQGTLVTVSSIQLTQSPSSLSASVGDRVTITCRASQGISAALAWYQQKPGKAPKLLIYAASSL
TSGVPSRFSGSGSGTDFTLTISSLQPEDFATYYCQQVYSFGGGTKVEIK
>P4.D37
EVQLVESGGGLVQPGRSLRLSCAASGFTFDDYAMHWVRQAPGKGLEWVSGISGNSGSIGYADSVKGRFTISRDNAKNSLYLQMNSL
RAEDTALYYCARTLDKANYFDYWGQGTLVTVSSIQLTQSPSSLSASVGDRVTITCRASQGISAALAWYQQKPGKAPKLLIYDASSL
DSGVPSRFSGSGSGTDFTLTISSLQPEDFATYYCQQVYSFGGGTKVEIK
>P4.D38
EVQLVESGGGLVQPGRSLRLSCAASGFTFDDYAMHWVRQAPGKGLEWVSGISSNSGSIGYADSVKGRFTISRDNAKNSLYLQMNSL
RAEDTALYYCARTIDMANYFDYWGQGTLVTVSSIQLTQSPSSLSASVGDRVTITCRASQGISAALAWYQQKPGKAPKLLIYAASSL
ESGVPSRFSGSGSGTDFTLTISSLQPEDFATYYCQQVYSFGGGTKVEIK
>P4.D39
EVQLVESGGGLVQPGRSLRLSCAASGFTFDDYAMHWVRQAPGKGLEWVSGISSNSGSIGYADSVKGRFTISRDNAKNSLYLQMNSL
RAEDTALYYCARTVDNANYFDYWGQGTLVTVSSIQLTQSPSSLSASVGDRVTITCRASQGISAALAWYQQKPGKAPKLLIYAASSL
DSGVPSRFSGSGSGTDFTLTISSLQPEDFATYYCQQVYSFGGGTKVEIK
>P4.D40
EVQLVESGGGLVQPGRSLRLSCAASGFTFDDYAMHWVRQAPGKGLEWVSGISSNSGSIGYADSVKGRFTISRDNAKNSLYLQMNSL
RAEDTALYYCARTLDKSNYFDYWGQGTLVTVSSIQLTQSPSSLSASVGDRVTITCRASQGISAALAWYQQKPGKAPKLLIYDASSL
HSGVPSRFSGSGSGTDFTLTISSLQPEDFATYYCQQVYSFGGGTKVEIK
>P4.D41
EVQLVESGGGLVQPGRSLRLSCAASGFTFDDYAMHWVRQAPGKGLEWVSGISSNSGSIGYADSVKGRFTISRDNAKNSLYLQMNSL
RAEDTALYYCARTLYSDNYFDYWGQGTLVTVSSIQLTQSPSSLSASVGDRVTITCRASQGISAALAWYQQKPGKAPKLLIYAASSL
DSGVPSRFSGSGSGTDFTLTISSLQPEDFATYYCQQVYSFGGGTKVEIK
>P4.D42
EVQLVESGGGLVQPGRSLRLSCAASGFTFDDYAMHWVRQAPGKGLEWVSGISSNSGSIGYADSVKGRFTISRDNAKNSLYLQMNSL
RAEDTALYYCARTLDKANYFDYWGQGTLVTVSSIQLTQSPSSLSASVGDRVTITCRASQGISAALAWYQQKPGKAPKLLIYAASSL
DSGVPSRFSGSGSGTDFTLTISSLQPEDFATYYCQQVYSFGGGTKVEIK
>P4.D43
EVQLVESGGGLVQPGRSLRLSCAASGFTFDDYAMHWVRQAPGKGLEWVSGISGNSGSIGYADSVKGRFTISRDNAKNSLYLQMNSL
RAEDTALYYCARTLDKDNYFDYWGQGTLVTVSSIQLTQSPSSLSASVGDRVTITCRASQGISAALAWYQQKPGKAPKLLIYAASSL
DSGVPSRFSGSGSGTDFTLTISSLQPEDFATYYCQQVYSFGGGTKVEIK
>P4.D44
EVQLVESGGGLVQPGRSLRLSCAASGFTFDDYAMHWVRQAPGKGLEWVSGISSNSGSIGYADSVKGRFTISRDNAKNSLYLQMNSL
RAEDTALYYCARTINKANYFDYWGQGTLVTVSSIQLTQSPSSLSASVGDRVTITCRASQGISAALAWYQQKPGKAPKLLIYAASSL
ESGVPSRFSGSGSGTDFTLTISSLQPEDFATYYCQQVYSFGGGTKVEIK
>P4.D45
EVQLVESGGGLVQPGRSLRLSCAASGFTFDDYAMHWVRQAPGKGLEWVSGISSNSGSIGYADSVKGRFTISRDNAKNSLYLQMNSL
RAEDTALYYCARTLDKSNYFDYWGQGTLVTVSSIQLTQSPSSLSASVGDRVTITCRASQGISAALAWYQQKPGKAPKLLIYDASSL
ESGVPSRFSGSGSGTDFTLTISSLQPEDFATYYCQQVYSFGGGTKVEIK

**>P5**
**QVQLQESGPGLVKPSETLSLTCAVYGGSFSGYYWSWIRQPPGKGLEWIGEINSGSTNYNPSLKSRITMSVDTSKNQFYLKLSSVTA**
**ADTAVYYCATLTGELDAFDVWGQGTLVTVSSSYELTQPLSVSVALGQAARITCGGNNLGYKSVWYQQKPGQAPVLVIYRDNNRPSG**
**IPERFSGSNSGNTATLTISRAQAGDEADYYCQSYDSSNVVFGSGTKVTVL**

>P5.D1
QVQLQESGPGLVKPSETLSLTCAVYGGSFSGYWWSWIRQPPGKGLEWIGQINHSGATQYNPSLKSRITMSVDTSKNQFYLKLSSVT
AADTAVYYCATLTGWLDAFDVWGQGTLVTVSSYELTQPLSVSVALGQAARITCGGNNLGYKSVHWYQQKPGQAPVLVIYRDNNRPS
GIPERFSGSNSGNTATLTISRAQAGDEADYYCQSYDGSNVVFGSGTKVTVL
>P5.D2
QVQLQESGPGLVKPSETLSLTCAVYGGSFSGYWWSWIRQPPGKGLEWIGQINHSGATMYNPSLKSRITMSVDTSKNQFYLKLSSVT
AADTAVYYCATLTGDLDAFDVWGQGTLVTVSSYELTQPLSVSVALGQAARITCGGNNLGYKSVHWYQQKPGQAPVLVIYRDNNRPS
GIPERFSGSNSGNTATLTISRAQAGDEADYYCQSYDGSNVVFGSGTKVTVL
>P5.D3
QVQLQESGPGLVKPSETLSLTCAVYGGSFSGYWWSWIRQPPGKGLEWIGMINHSGATMYNPSLKSRITMSVDTSKNQFYLKLSSVT
AADTAVYYCATLTGDLDAFDVWGQGTLVTVSSYELTQPLSVSVALGQAARITCGGNNLGYKSVHWYQQKPGQAPVLVIYRDNNRPS
GIPERFSGSNSGNTATLTISRAQAGDEADYYCQSYDGSNVVFGSGTKVTVL
>P5.D4
QVQLQESGPGLVKPSETLSLTCAVYGGSFSGYWWSWIRQPPGKGLEWIGQINHSGATQYNPSLKSRITMSVDTSKNQFYLKLSSVT
AADTAVYYCATLTGDLDAFDVWGQGTLVTVSSYELTQPLSVSVALGQAARITCGGNNLGYKSVHWYQQKPGQAPVLVIYRDNNRPS
GIPERFSGSNSGNTATLTISRAQAGDEADYYCQSYDGSNVVFGSGTKVTVL
>P5.D5
QVQLQESGPGLVKPSETLSLTCAVYGGSFSGYWWSWIRQPPGKGLEWIGQINHSGAVMYNPSLKSRITMSVDTSKNQFYLKLSSVT
AADTAVYYCATLTGDLDAFDVWGQGTLVTVSSYELTQPLSVSVALGQAARITCGGNNLGYKSVHWYQQKPGQAPVLVIYRDNNRPS
GIPERFSGSNSGNTATLTISRAQAGDEADYYCQSYDGSNVVFGSGTKVTVL
>P5.D6
QVQLQESGPGLVKPSETLSLTCAVYGGSFSGYWWSWIRQPPGKGLEWIGAINHSGATKYNPSLKSRITMSVDTSKNQFYLKLSSVT
AADTAVYYCATLTGDLDAFDVWGQGTLVTVSSYELTQPLSVSVALGQAARITCGGNNLGYKSVHWYQQKPGQAPVLVIYRDNNRPS
GIPERFSGSNSGNTATLTISRAQAGDEADYYCQSYDGSNVVFGSGTKVTVL
>P5.D7
QVQLQESGPGLVKPSETLSLTCAVYGGSFSGYWWSWIRQPPGKGLEWIGQINHSGATAYNPSLKSRITMSVDTSKNQFYLKLSSVT
AADTAVYYCATLTGDLDAFDVWGQGTLVTVSSYELTQPLSVSVALGQAARITCGGNNLGYKSVHWYQQKPGQAPVLVIYRDNNRPS
GIPERFSGSNSGNTATLTISRAQAGDEADYYCQSYDGSNVVFGSGTKVTVL

#### S3. Average number of mutations by affinity maturation for each prototype sequence

| P1 | P2 | P3 | P4 | P5 |
|---|---|---|---|---|
| 5.42 | 4.54 | 3.56 | 4.25 | 4.57 |

#### S4. Alanine scanning using PyRosetta

An available script from PyRosetta[2] scripts repository was used for alanine scanning.

https://graylab.jhu.edu/pyrosetta/downloads/scripts/demo/D090_Ala_scan.py

#### Results of alanine scanning for top 5 selected affinity maturation designs

| P1D1 | | P1D2 | | P4D1 | | P3D1 | | P3D3 | |
|---|---|---|---|---|---|---|---|---|---|
| Interface residue | Loss in binding energy kcal/mol | Interface residue | Loss in binding energy kcal/mol | Interface residue | Loss in binding energy kcal/mol | Interface residue | Loss in binding energy kcal/mol | Interface residue | Loss in binding energy kcal/mol |
| G62 | 9.69 | I114 | 16.01 | G447 | 3.735 | W64 | 2.298 | N57 | 0.99 |
| F109 | 3.041 | G62 | 9.73 | G446 | 2.2 | N57 | 0.648 | G58 | 0.78 |
| G63 | 2.128 | F109 | 2.763 | A56 | 0.573 | F107 | 0.567 | F107 | 0.33 |
| L65 | 1.26 | T65 | 1.42 | Y489 | 0.209 | S29 | 0.14 | S29 | 0.22 |
| W353 | 1.031 | Y66 | 1.06 | D35 | 0.083 | S108 | 0.117 | D38 | 0.182 |
| I56 | 0.938 | I56 | 0.94 | S494 | 0.021 | D38 | 0.11 | R357 | 0.099 |
| L38 | 0.725 | S57 | 0.706 | C488 | 0.02 | I29 | 0.079 | Q106 | 0.03 |
| W59 | 0.706 | W353 | 0.51 | G28 | 0.01 | Q106 | 0.016 | I29 | 0.029 |
| S57 | 0.685 | L38 | 0.507 | A25 | 0.005 | G446 | 0.013 | I2 | 0.027 |
| Y66 | 0.53 | G63 | 0.5 | A348 | 0.005 | C336 | 0.008 | S108 | 0.021 |
| N38 | 0.386 | Y489 | 0.467 | T85 | 0.004 | | | C336 | 0.02 |
| N487 | 0.34 | N487 | 0.372 | W353 | 0.004 | | | R36 | 0.001 |
| C488 | 0.333 | C488 | 0.27 | L67 | 0.003 | | | | |
| R80 | 0.059 | I78 | 0.18 | A352 | 0.003 | | | | |
| A64 | 0.052 | T77 | 0.136 | F347 | 0.002 | | | | |
| T345 | 0.037 | N354 | 0.113 | | | | | | |
| I78 | 0.023 | N38 | 0.108 | | | | | | |
| I2 | 0.01 | P337 | 0.057 | | | | | | |
| | | R80 | 0.052 | | | | | | |
| | | R346 | 0.031 | | | | | | |

### S5. Molecular dynamics of antibody design P1.D1 in complex with SARS-CoV2-RBD

All atom MD simulations were performed using Desmond[3] release of Schrodinger 2019.4 release[8]. The OPLS3[4] force field was used with TIP3P[5] water model. NaCl ions were added at 0.15M concentration to neutralize the solvated system. Equilibration of the system was carried using the default relaxation protocol in Desmond. The protocol started with an energy minimization followed by a 12 ps long NVT simulation at 10 K followed by a 12ps long NPT simulation – all with restraints on the solute. Then the system was equilibrated at 310K in the NPT ensemble for 24 ps using 2 fs time step without any restraints. The production simulation was 50ns long with 2fs time step at 310 K in the NPT ensemble. The trajectories were analyzed using VMD[6] analysis plugin. RMSD analysis was done using *RMSD Visualizer plugin*. Hydrogen bonds were calculated using *Hydrogen Bonds plugin.* Interaction energy was calculated using *interaction_energy.py* module of Desmond package.

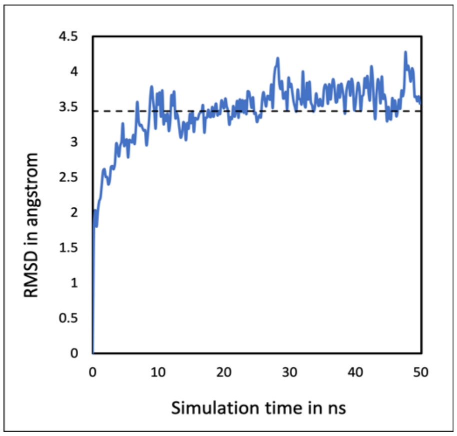


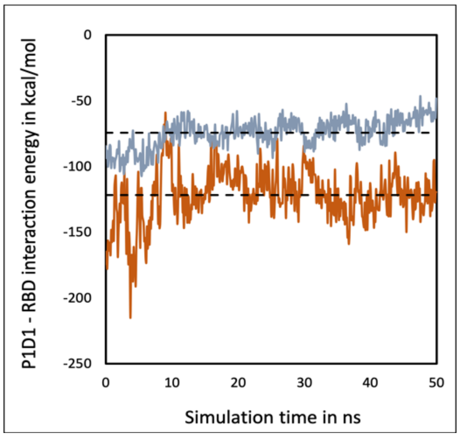


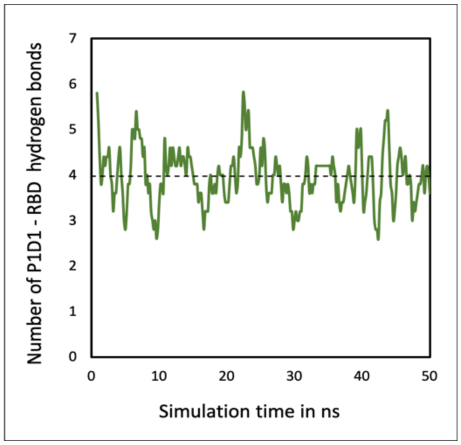


**Figure S5. a). Plot showing all atom-RMSD after aligning upon the initial structure at 0 ns b). Plot showing interaction energy between antibody and antigen – in Gray is the van-der-Waals energy and in Red is the Electrostatic energy c). Plot showing number of Hydrogen bonds at the interface of antibody-antigen calculated using VMD plugin with default parameters. In each plot, dotted lines show the average value.**

### S6. Rosetta affinity maturation protocol

The protocol was implemented using RosettaScripts[7] with the following xml script. (A sample flags file and resfile used are given below)

### XML script: (dock_design.xml)

```
<ROSETTASCRIPTS>
	<SCOREFXNS>
		<ScoreFunction name="r15" weights="ref2015" />
	</SCOREFXNS>
	<TASKOPERATIONS>
		<InitializeFromCommandline name="ifcl"/>
		<RestrictToRepacking name="rtr" />
		Design and repack residues based on resfile
		<ReadResfile name="rrf" filename="%%resfile%%"/>
	</TASKOPERATIONS>
	<FILTERS>
	</FILTERS>
	<MOVERS>
		MINIMIZATION MOVERS

		Relax is taking too long!
		<MinMover name="minimize" jump="all" chi="True" bb="True" scorefxn="r15" >
		</MinMover>

		<Backrub name="backrub_motion" pivot_residues="%%pivots%%" />
		<GenericMonteCarlo name="backrub" mover_name="backrub_motion"
scorefxn_name="REF2015" trials="500" temperature="1.0" recover_low="1" />

		Single cycle of FastRelax to minimize backbone of docking partners
		<FastRelax name="relax" scorefxn="REF2015" repeats="1" task_operations="ifcl,rtr"
/>

		DOCKING MOVERS
		<DockingProtocol name="full_dock" docking_score_high="r15" dock_min="1"
ignore_default_docking_task="0" task_operations="ifcl,rtr,rrf" partners="%%partners%%"/>
		<DockingProtocol name="hires_dock" docking_score_high="r15"
docking_local_refine="1" dock_min="1" ignore_default_docking_task="0"
task_operations="ifcl,rtr,rrf" partners="%%partners%%"/>

		DESIGN MOVER
		<PackRotamersMover name="design" scorefxn="REF2015" task_operations="ifcl,rrf" />
		<PackRotamersMover name="packonly" scorefxn="REF2015" task_operations="ifcl,rtr"
/>


		INTERFACE METRICS
		<InterfaceAnalyzerMover name="analyze" scorefxn="REF2015" packstat="1"
pack_input="0" pack_separated="1" interface="%%partners%%" />

		No idea why MPI does not output all structures
		Trying this in desperation
		<DumpPdb name="save" fname="dump.pdb" tag_time="true" scorefxn="REF2015" />

	</MOVERS>
	<APPLY_TO_POSE>
	</APPLY_TO_POSE>
	<PROTOCOLS>
		Run docking protocol
		<Add mover="hires_dock"/>

		Backrub
		<Add mover="backrub" />
		Design
		<Add mover="design" />
		pack
		<Add mover="packonly" />
```

```
                Backrub
                <Add mover="backrub" />
                Design
                <Add mover="design" />
                pack
                <Add mover="packonly" />

                Backrub
                <Add mover="backrub" />
                Design
                <Add mover="design" />
                pack
                <Add mover="packonly" />

                Hires-dock
                <Add mover="hires_dock"/>

                Minimize
                <Add mover="minimize" />

                Dump
                <Add mover="save" />
                Calculate interface metrics for the final sequence
                Add mover="analyze" />

        </PROTOCOLS>
        <OUTPUT scorefxn="REF2015" />
</ROSETTASCRIPTS>
```

### Sample flags file: (P5.flags)

```
# defaults
-ex1
-ex2
-ex2aro
-use_input_sc
-ignore_zero_occupancy false

# 0 -> input pdb prepared file
-in:file:s P5.pdb

# 1 -> script file
-parser:protocol dock_score.xml

# 2 -> variable for partners
-parser:script_vars partners=A_HL

# 3 -> nstruct
-nstruct 1

# 4 -> output suffix
-out:suffix _scorenow

# 5 -> variable for resfile
-parser:script_vars resfile=P5.resfile

# 6 -> pivot residues
-parser:script_vars
pivots=2H,3H,4H,5H,6H,7H,8H,9H,11H,12H,13H,14H,15H,16H,17H,18H,19H,20H,21H,22H,23H,24H
,25H,26H,27H,28H,29H,30H,35H,36H,37H,38H,39H,40H,41H,42H,43H,44H,45H,46H,47H,48H,49H,5
0H,51H,52H,53H,54H,55H,56H,57H,58H,59H,62H,63H,64H,65H,66H,67H,68H,69H,70H,71H,72H,74H
,75H,76H,77H,78H,79H,80H,81H,82H,83H,84H,85H,86H,87H,88H,89H,90H,91H,92H,93H,94H,95H,9
6H,97H,98H,99H,100H,101H,102H,103H,104H,105H,106H,107H,108H,109H,110H,111H,112H,113H,1
14H,115H,116H,117H,118H,119H,120H,121H,122H,123H,124H,125H,126H,127H,2L,3L,4L,5L,6L,7L
,8L,9L,10L,11L,12L,13L,14L,15L,16L,17L,18L,19L,20L,21L,22L,23L,24L,25L,26L,27L,28L,29L
,36L,37L,38L,39L,40L,41L,42L,43L,44L,45L,46L,47L,48L,49L,50L,51L,52L,53L,54L,55L,56L,5
7L,65L,66L,67L,68L,69L,70L,71L,72L,74L,75L,76L,77L,78L,79L,80L,83L,84L,85L,86L,87L,88L
```

```
,89L,90L,91L,92L,93L,94L,95L,96L,97L,98L,99L,100L,101L,102L,103L,104L,105L,106L,107L,1
08L,109L,114L,115L,116L,117L,118L,119L,120L,121L,122L,123L,124L,125L,126L,
```

### Sample resfile: (P5.resfile)

```
NATRO
start
37 H ALLAA
38 H ALLAA
57 H ALLAA
59 H ALLAA
62 H ALLAA
63 H ALLAA
64 H ALLAA
65 H ALLAA
66 H ALLAA
108 K ALLAA
109 K ALLAA
110 H ALLAA
114 K ALLAA
116 K ALLAA
346 A NATAA
347 A NATAA
348 A NATAA
351 A NATAA
352 A NATAA
354 A NATAA
355 A NATAA
```

### Sample command:

```
rosetta_scipts.mpi.linuxgccrelease @P5.flags
```

### S7. Binding mode of CR3022 compared with that of P1.D1

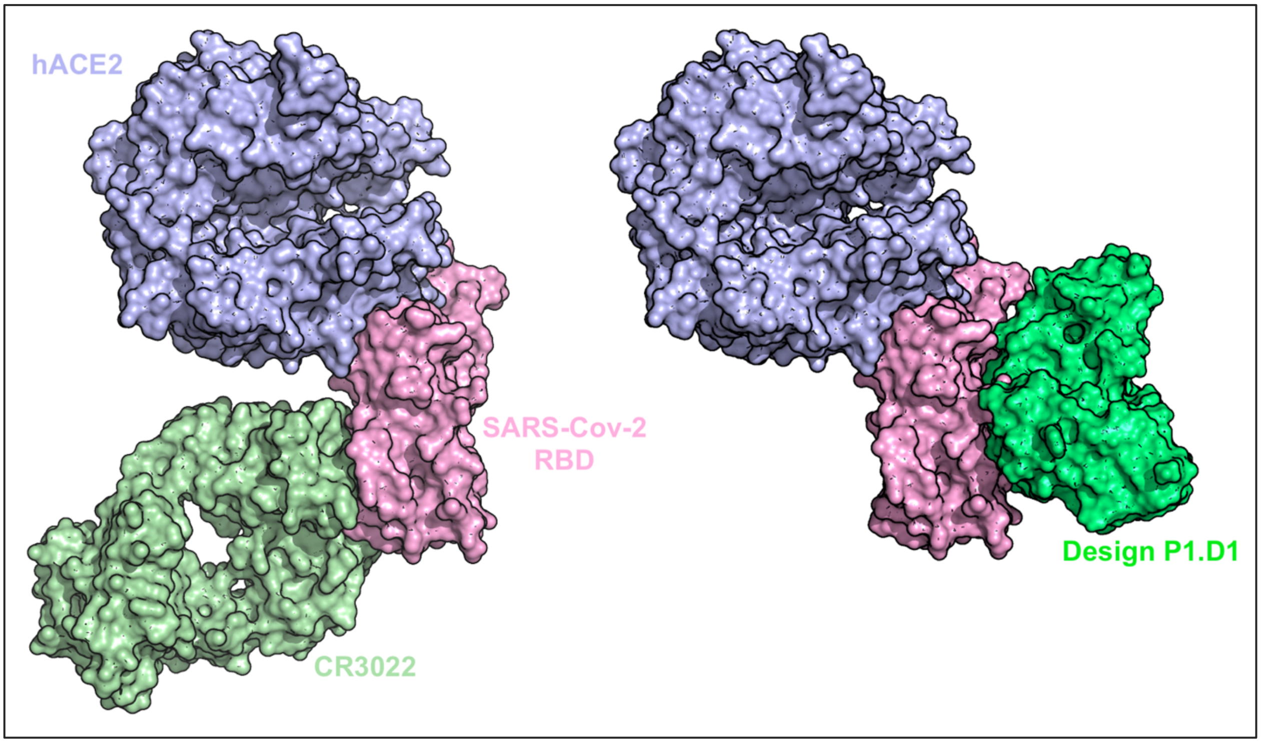


**Fig. S7.** (Left) CR3022 bound to RBD (PDB-id: 6w41) superposed upon ACE2 bound to RBD (PDB-id: 6lzg). (Right) P1.D1 design bound to RBD superposed upon ACE2 bound to RBD. Both CR3022 and P1.D1 are bound to two different epitopes on the RBD which are both different from the ACE2 binding site.

**S8. Binding location of P1.D1 in the full spike protein (open confirmation) compared with that of hACE2**

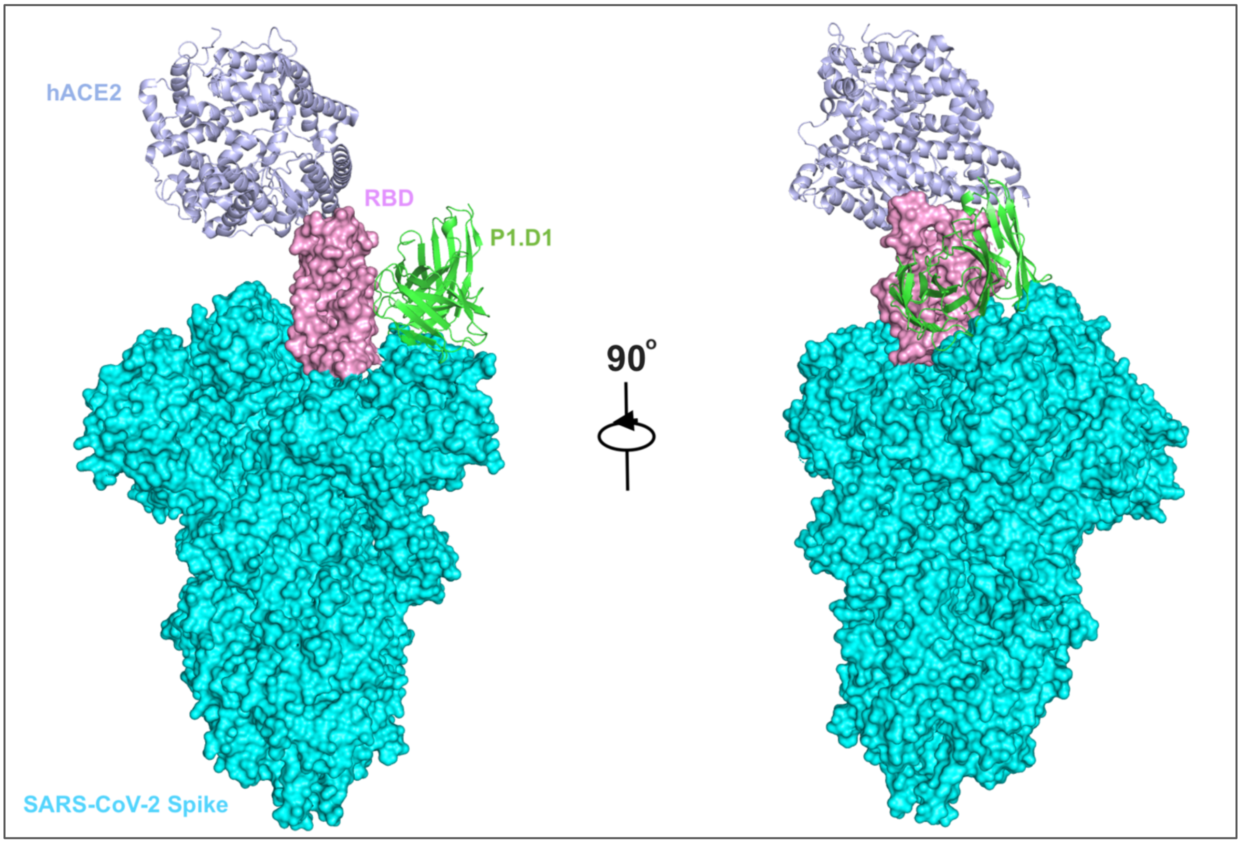


**Fig S8.** Figure showing the P1.D1-RBD complex and hACE2-RBD complex (PDB:- 6LZG) superimposed on the SARS-CoV-2 spike protein open confirmation (PDB:- 6VYB) illustrating that P1.D1 binds to an epitope on the RBD exposed in the open confirmation. The spike protein is shown in space-filling mode, hACE2 and P1.D1 are shown as cartoons.
